# Frictiotaxis underlies adhesion-independent durotaxis

**DOI:** 10.1101/2023.06.01.543217

**Authors:** Adam Shellard, Kai Weißenbruch, Peter A. E. Hampshire, Namid R. Stillman, Christina L. Dix, Richard Thorogate, Albane Imbert, Guillaume Charras, Ricard Alert, Roberto Mayor

## Abstract

Cells move directionally along gradients of substrate stiffness, a process called durotaxis. The current consensus is that durotaxis relies on cell-substrate focal adhesions to sense stiffness and transmit forces that drive directed motion. Therefore, focal adhesion-independent durotaxis is thought to be impossible. Here, we show that confined cells can perform durotaxis despite lacking strong or specific adhesions. This durotactic migration depends on asymmetric myosin distribution and actomyosin retrograde flow. We show that the mechanism of this adhesion-independent durotaxis is that stiffer substrates offer higher friction. We propose a physical model that predicts that non-adherent cells polarise and migrate towards regions of higher friction – a process that we call frictiotaxis. We demonstrate frictiotaxis in experiments by showing that cells migrate up a friction gradient even when stiffness is uniform. Our results broaden the potential of durotaxis to guide any cell that contacts a substrate and reveal a new mode of directed migration based on friction, with implications for immune and cancer cells, which commonly move with non-specific interactions.

---

The ability of cells to migrate following environmental gradients underlies many aspects of development, homeostasis and disease^1,2^. Cells follow gradients in the stiffness of their substrate, a process called durotaxis that has been demonstrated in multiple cell types *in vitro* and *in vivo*^3–7^. The prevailing mechanistic view of durotaxis involves the cells’ actomyosin machinery producing contractile forces that pull on the underlying substrate through focal adhesions. These pulling forces then bias cell motion in one direction, typically toward the stiffer substrate^5,8–10^. These physical models are supported by experimental evidence, and an essential component of all of them is strong cell-substrate adhesions. It is thus believed that cells lacking adhesions must be incapable of sensing stiffness gradients. Whether cells lacking cell-matrix adhesions can respond to mechanical gradients is widely acknowledged as a vital question^11–14^ that has been unaddressed. Its importance is highlighted by the fact that adhesion-independent motility is a fundamental mode of migration, classically exhibited by various cancer and immune cells, but which can be triggered in practically any cell type by providing 3D confinement^15,16^.

## Microchannels with tuneable stiffness

Part of the challenge in addressing whether cells migrating without adhesions are capable of responding to stiffness gradients, or indeed the role of substrate stiffness in the context of 3D adhesion-independent motility in general, lies in the technical challenge of fabricating confined cellular environments of tuneable stiffness that are necessary to study adhesion-independent migration^12^. Adhesion-independent motility has commonly been studied using the ‘under agarose assay’ in which cells emigrate from a free region into confinement between agarose and glass. However, this method is unable to differentiate between the effect of confinement and the effect of compression since cells must deform their substrate to move. A newer method uses microchannels fabricated with polydimethylsiloxane (PDMS). However, this material exhibits stiffness in the MPa range, far from physiologically relevant levels of most tissues, and offers little ability to tune rigidity. We developed a novel and easy-to-use method in which mobile and deformable cells spontaneously migrate into and within agarose-surrounded preformed microchannels (Fig. 1a). Fabricating microchannels with agarose offers the potential to investigate cells within substrates of physiologically relevant stiffnesses^17^. The dimensions of the channel are determined by a PDMS mould that provides reliable confinement irrespective of microchannel dimensions (Fig. 1b,c and Extended Data Fig. 1a-c) and is unaffected by the stiffness of the substrate (Fig. 1e,g and Extended Data Fig. 1d,e). The stiffness of the microchannel is tuned by the concentration of agarose (Fig. 1d). Fluorospheres did not show directed motion within microchannels (Extended Data Fig. 1f,g), confirming that pressure-driven fluid flow was not a factor in this setup. Importantly, stiffness gradients can be created by combining solutions with different agarose concentrations together during microchannel assembly (Fig. 1h,i).

**Figure 1.**
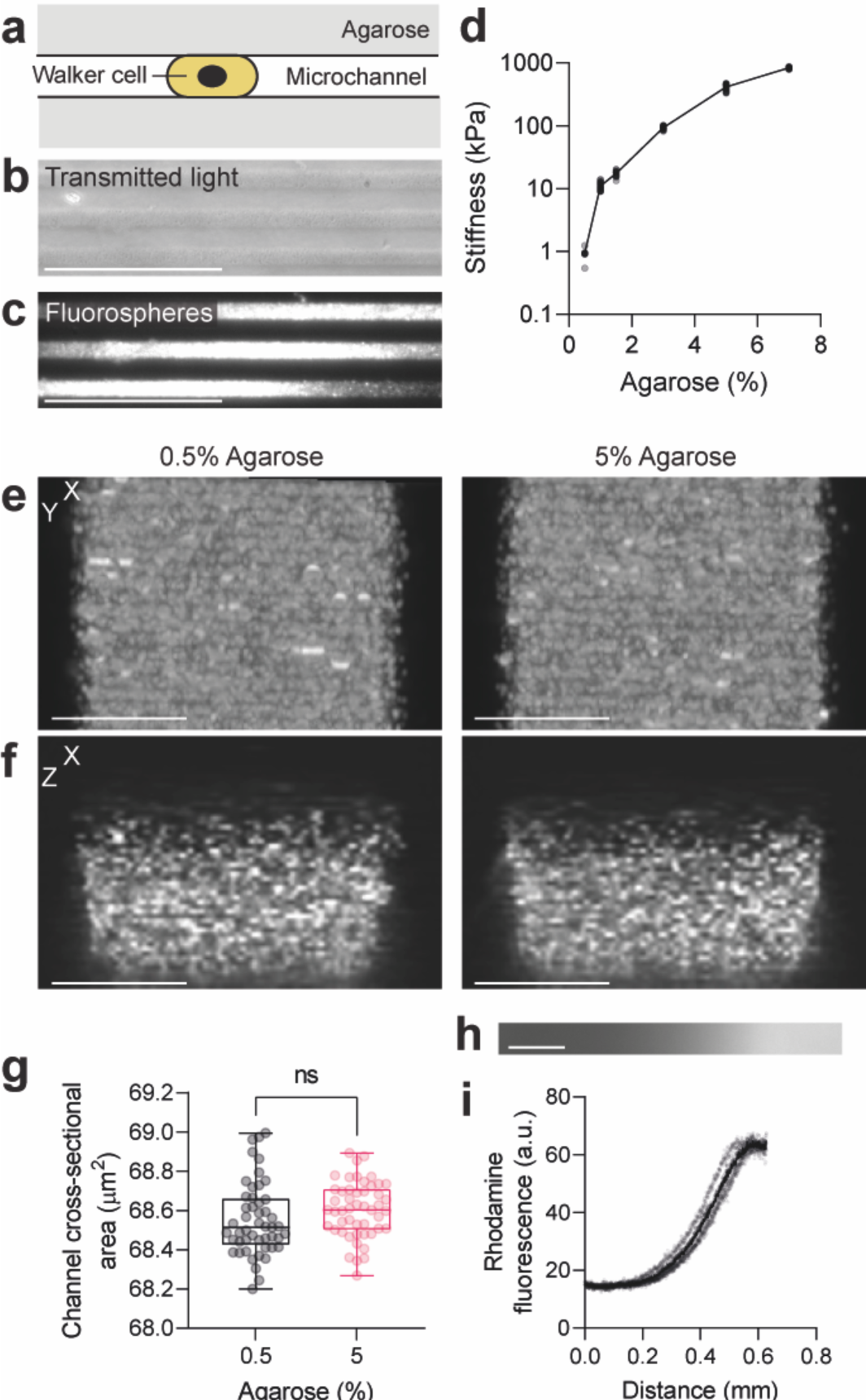
A microchannel system with tuneable stiffness and dimensions. **a**, Diagram of the microchannel assay. Microchannel dimensions determine the geometry of the setup and the agarose surrounding the cell determines substrate stiffness. **b-c**, Agarose channels (**b**) filled with 0.2 µm fluorospheres (**c**). Scale bar, 100 µm. **d**, Stiffness measurements (mean ± s.d.) by nanoindentation of gels of different agarose concentration. Grey dots represent individual data points, black dots represent mean. *n* = 25 cells. **e-g**, Agarose microchannels (**e-f**; 0.5%, left; 5%, right) filled with 0.2 µm fluorospheres and imaged from below (**e**, maximum projection) and side-on (**f**). Channel cross-sectional area quantified (**g**). Scale bar, 5 µm. For **g**, n *=* 49 channels; unpaired two-tailed t test; ns, *P*>0.05 (exact: *P*=0.2229). **h-i**, A stiffness gradient visualized by immersing rhodamine dextran dye into one of the agarose solutions prior to subsequent diffusion and solidification (**h**) and quantification of the dye fluorescence over the gradient axis (**i**). Scale bar, 100 µm. Grey dots represent individual data points, black dots represent mean. *n* = 5 gels.

To test our setup as a valid means of assaying confined non-adherent cellular motility, we used a non-adherent subline of Walker 256 carcinosarcoma (henceforth Walker) cells as a well-validated model of a cell type that moves without using specific substrate adhesions and without focal adhesions^18,19^. Although Walker cells are able to attach to fibronectin, they are completely non-adhesive on agarose or on glass coated with PLL-PEG (Fig. 2a,b, Extended Data Fig. 2). Furthermore, Walker cells introduced into the agarose microchannel setup exhibited classical amoeboid motion characterised by bleb-based rather than lamellipodium-based motility (Fig. 2c,d and Supplementary Video 1), fast migration (Fig. 2e) and lack of focal adhesions (Fig. 2f,g). Together, these observations indicate that Walker cells are non-adhesive in our agarose microchannel setup.

**Figure 2.**
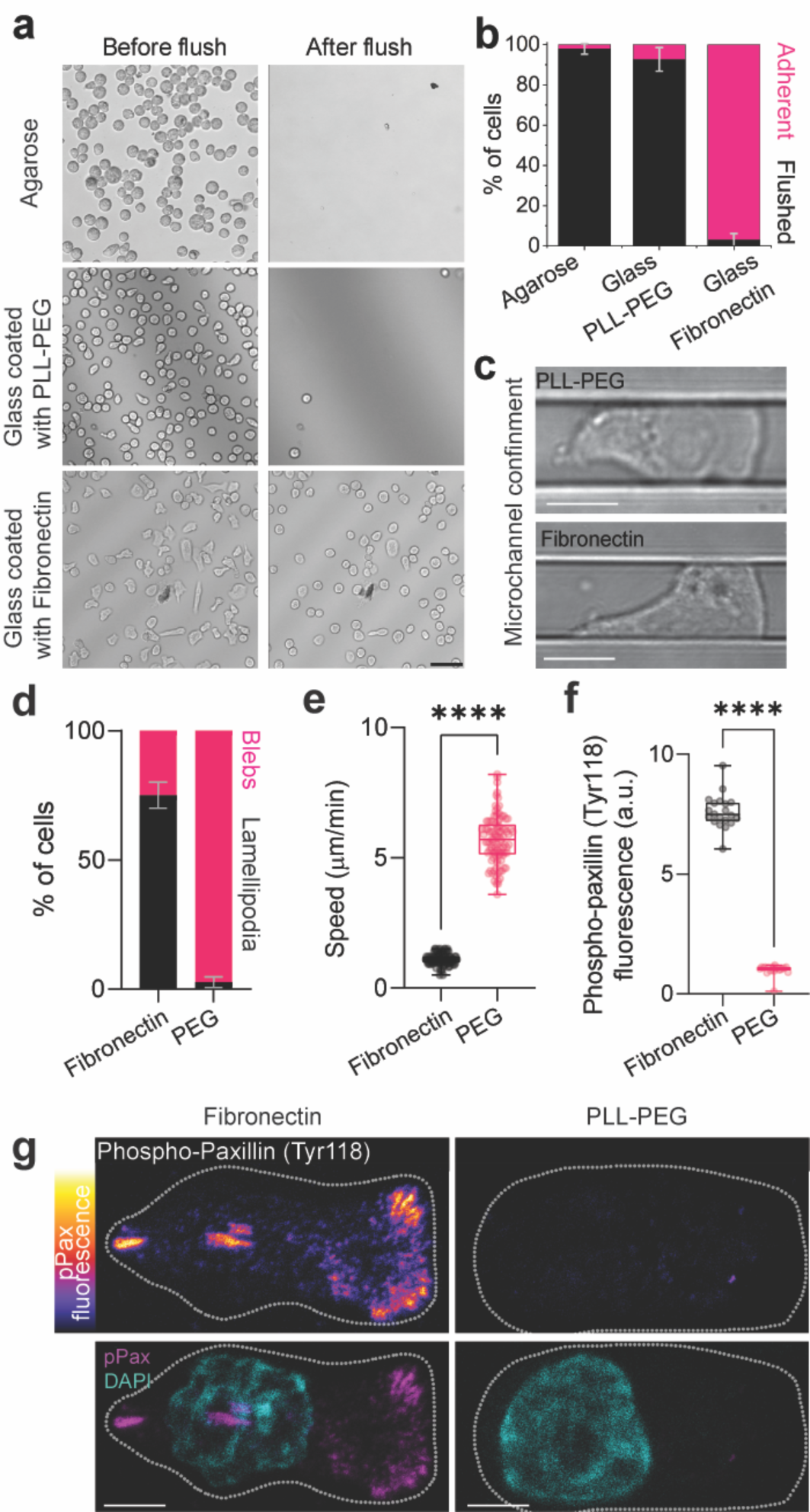
Walker cells exhibit adhesion-independent amoeboid migration in the microchannel system. **a**, Images of Walker cells on agarose, glass coated with PLL-PEG, and glass coated with fibronectin, before and after flushing the substrate with culture medium. Scale bar, 50 µm. **b**, Quantification of adherent and flushed fraction of cells after flushing the respective substrates. Bars represent mean ± s.d.; *N =* 3 experimental repeats. **c**, Example images of cells in non-adhesive (PLL-PEG, top) or adhesive (fibronectin, bottom) agarose microchannels. Scale bar, 10 µm. **d**, Quantification of lamellipodia and blebs by cells in adhesive (fibronectin) and non-adhesive (PEG) conditions. Bars represent mean ± s.d.; *N =* 3 experimental repeats. **e,** Speed of cells migrating within agarose microchannels with fibronectin or PEG coating. **f-g**, Immunostaining against phospho-paxillin (Tyr118) in Walker cells within agarose channels (**g**) and quantification (**f**) in which the underlying glass is coated with either PLL-PEG or fibronectin. Dotted white line represents the cell outline. Scale bar, 5 µm. *n* = 100 cells (**e, f**); two-tailed Mann-Whitney test; \*\*\*\**P*≤0.0001.

## Adhesion-independent durotaxis

To directly address whether cells are capable of undergoing durotaxis in an adhesion-independent manner, agarose microchannels were fabricated to have either uniform stiffness or a stiffness gradient (Fig. 3a). We tracked migratory cells that entered regions of the microchannel which exhibited either uniform or graded stiffness. Cells entering substrate of uniform stiffness had a moderate tendency to repolarise and move back, whereas cells entering a stiffness gradient were characterised by much higher persistence as they migrated toward the stiff substrate (Fig. 3b-e and Supplementary Video 2). In addition, cells exhibited a higher migration speed at higher stiffness values (Fig. 3f and Extended Data Fig. 3). Like Walker cells, HL60 neutrophil-like cells, a cell type with a well-described amoeboid behaviour^20^, also exhibited preferred motion toward stiffer substrate (Extended Data Fig. 4). Together, these results reveal adhesion-independent durotaxis may be a generalisable phenomenon.

**Figure 3.**
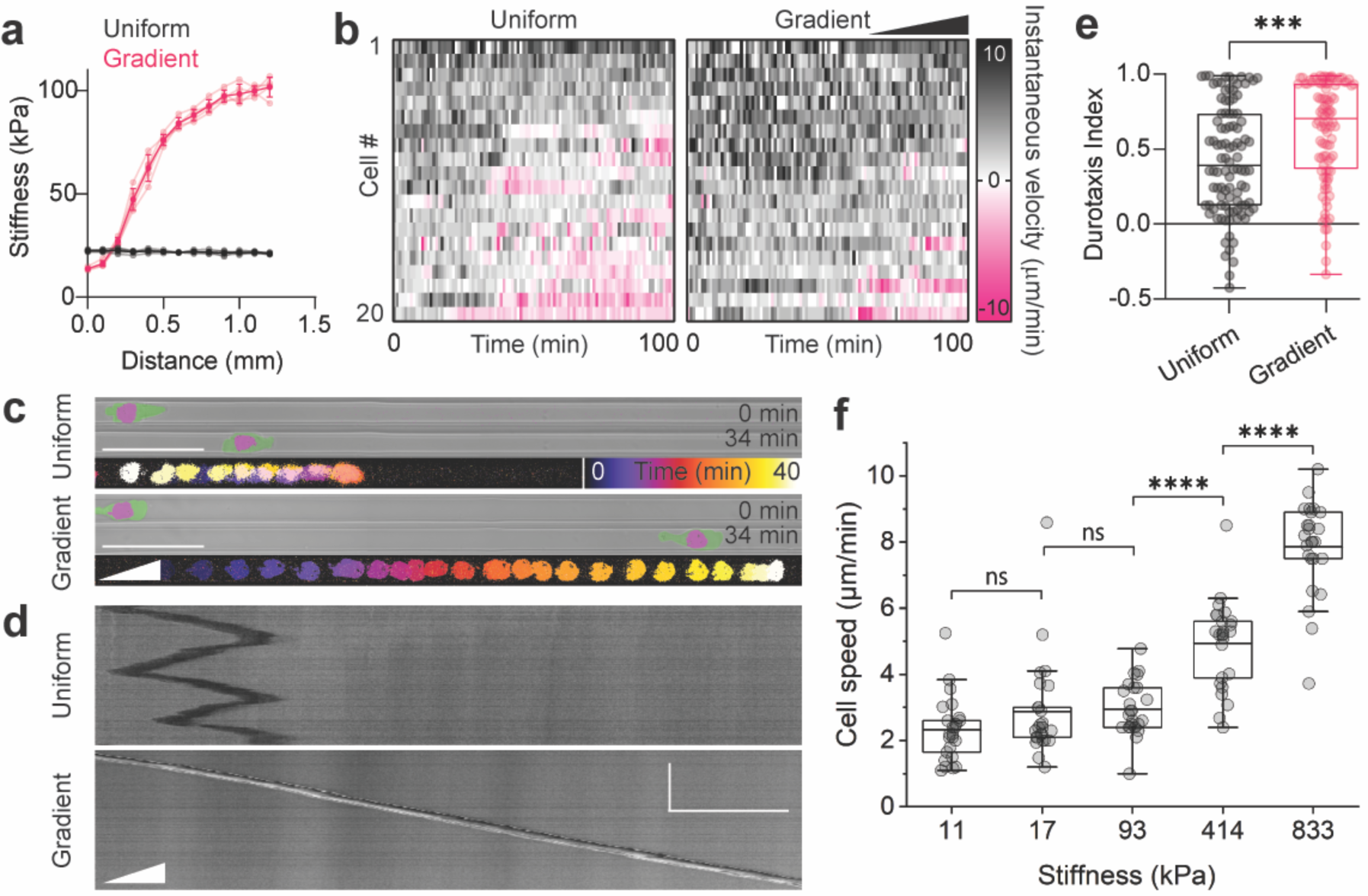
Adhesion-independent durotaxis of Walker cells. **a**, Stiffness measurements (mean ± s.d.) from nanoindentation of uniform (black) and gradient (pink) microchannels. Opaque dots and lines represent mean; translucent dots and lines represent raw data. *n* = 5 gels each. **b**, Heat maps of the instantaneous velocity of representative cells migrating within microchannels of uniform or graded stiffness. *n* = 20 representative cells each. Each row represents a different cell. The *x*-axis represents time, with each box representing 1 min for a total of 100 min. Each heat map is ordered such that cells migrating forward (black) are at the top, and cells migrating backwards (pink) are at the bottom. **c**, Example pictures of cells at an earlier (top panels) and later (middle panels) time point, and temporal colour-coded projected tracks (bottom panels). Cells are tracked by a nuclear marker and membrane is pseudocoloured. Scale bar, 50 µm. **d**, Kymographs of the cells shown in (**c**). Scale bar, 50 µm (horizontal), 50 min (vertical). **e**, Quantification of durotaxis. *n* = 100 cells; two-tailed Mann-Whitney test; \*\*\**P*≤0.001 (exact: *P* = 0.0003). **f**, Quantification of migration speed in microchannels of different stiffness regimes. *n* = 25 cells for each stiffness regime; two-tailed Mann-Whitney test; ns, *P*>0.05 (11 versus 17: *P* = 0.23258; 17 versus 93: *P* = 0.09795), \*\*\*\**P*≤0.0001.

## Ameboid durotaxis depends on actomyosin flow

It has previously been described that amoeboid migration on substrates of uniform stiffness depends on actomyosin retrograde flow^18^. We therefore asked whether actomyosin retrograde flow was also observed in our cells undergoing adhesion-independent durotaxis. Walker cells expressing myosin-GFP were placed in the agarose microchannels with either uniform stiffness or a stiffness gradient, followed by time-lapse imaging. Cells exhibited an accumulation of myosin at the rear and a clear myosin retrograde flow (Fig. 4a,b, Extended Data Fig. 5a-c and Supplementary Video 4&5). Upon cell repolarisation, the direction of retrograde actomyosin flow was reversed, leading to accumulation of actomyosin at the new cell rear (Fig. 4b and Supplementary Video 4). A clear positive correlation between actomyosin flow and cell speed was observed (Fig. 4c and Extended Data Fig 5c). To determine whether myosin was required for adhesion-independent durotaxis, myosin activity was reduced by treating cells with the ROCK inhibitor Y-27632. A clear loss in the rear accumulation of myosin (Fig. 4d), retrograde flow (Extended Data Fig 5d), bleb formation (Fig. 4g), and impairment of migration was observed in treated cells compared with control cells (Fig. 4e,f,h and Supplementary Video 6). These observations show that rear accumulation of myosin dependent on actomyosin retrograde flow is required for adhesion-independent durotaxis.

**Figure 4.**
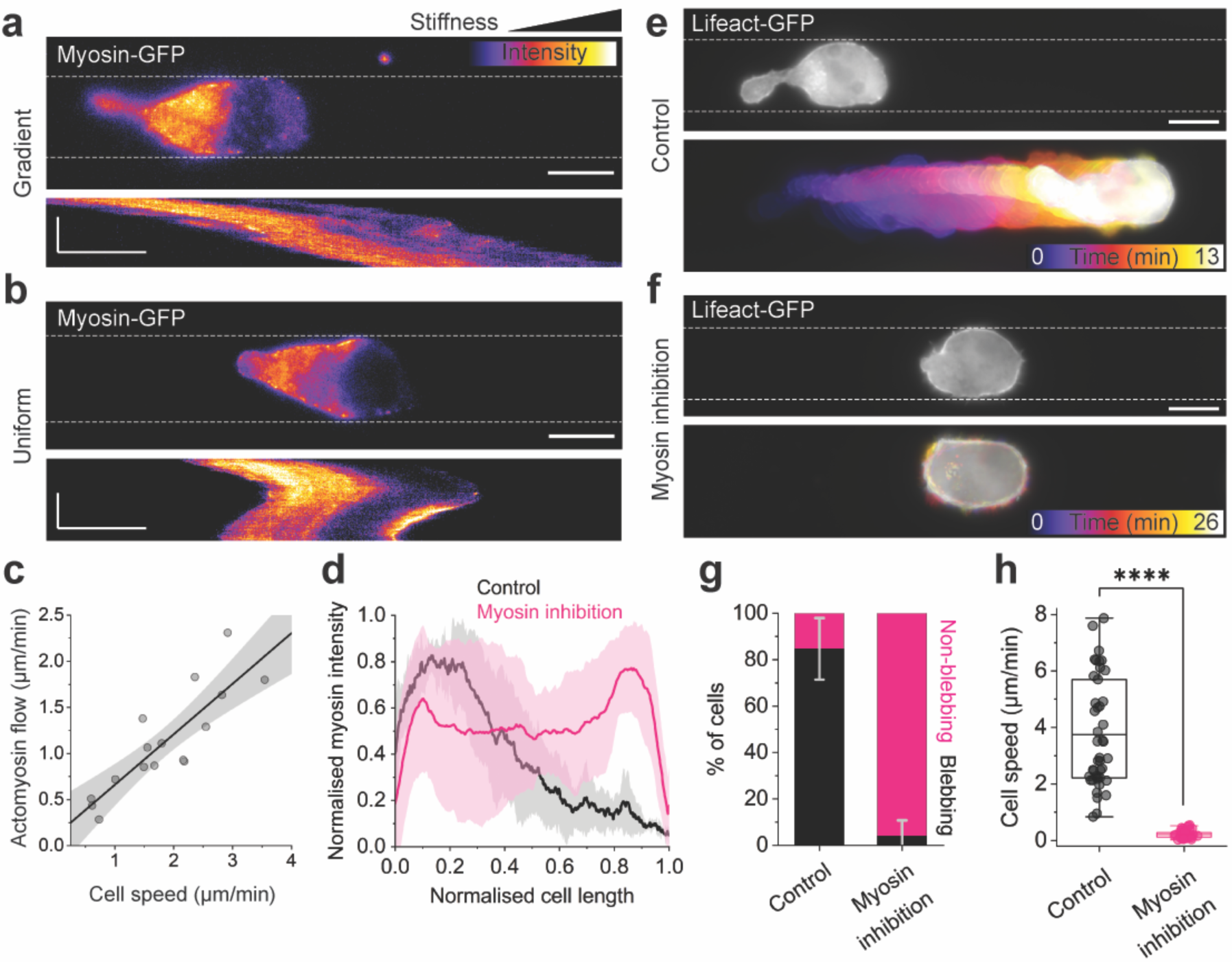
Adhesion-independent durotaxis depends on retrograde actomyosin flow. **a**-**b**, Still images and corresponding kymographs from timelapse records of Myosin-GPF-expressing Walker cells, migrating in microchannels with stiffness gradient (**a**) or uniform stiffness (**b**). Fluorescence signal was colour-coded, dotted lines represent channel boundaries. Scale bars, 10 µm in still images, 5 µm (horizontal) and 10 min (vertical) in kymographs. **c**, Correlation of actomyosin flow and migration speed. Coefficient of determination (*R*^2^) = 0.73068. Grey areas depict upper and lower 95% confidence limits. Dots represent mean values of individual cells migrating over 400 s. Data points of *n* = 8 cells in stiffness gradients or uniform stiffness were picked randomly from *N* = 3 independent experiments each and pooled. **d**, Intensity profiles of Myosin-GFP signals along the front-rear axis of Walker cells in the absence (DMSO control, black line) or presence of 30 µM Y-27632 (Myosin inhibition, purple line). Myosin-GFP intensity and cell length were normalised for comparison. Solid lines plus transparent areas depict mean ± s.d. of *n* = 6 cells each, from *N*=3 independent experiments. **e**-**f**, Still images and corresponding colour-coded timelapse records of Lifeact-GPF-expressing Walker cells in microchannels of uniform stiffness; in the absence (**e**) or presence of 30µM Y-27632 (**f**). Dotted lines represent channel boundaries. Scale bars, 10 µm. **g**, Quantification of blebbing and non-blebbing fractions of cells under control (DMSO) or myosin inhibiting (30 µM Y-27632) conditions. Bars represent mean ± s.d. from *N =* 3 independent experiments. **h**, Quantification of migration speed in microchannels of uniform stiffness. *n* = 43 cells for control (DMSO) and *n* = 47 cells for myosin inhibition (30 µM Y-27632); two-tailed Mann-Whitney test; \*\*\*\**P*≤0.0001.

## An active gel model of amoeboid migration predicts frictiotaxis

Given that amoeboid cells lack focal adhesions, it was unclear how these cells were able to durotax. We hypothesised that regions of higher stiffness may offer higher friction, which arises from non-specific molecular interactions between the cell membrane and the channel walls^18^. We also hypothesised that, in the absence of active forces pulling the substrate, this passive friction would cause amoeboid cells to undergo durotaxis.

We investigated this hypothesis through a physical model of amoeboid motility which treats the actomyosin network in the cell cortex as an active gel^13,15,16,18,21–25^. In this model, myosin-generated contractility triggers an instability whereby the gel concentrates towards one side, which becomes the cell rear (Fig. 5a). This concentration profile drives retrograde actomyosin flow, which propels the cell forward. In the absence of external gradients, symmetry is broken spontaneously. The cell can thus move either left or right (Fig. 5a), consistent with our experimental observations of repolarisation events in uniform channels (Fig. 3c,d).

**Figure 5.**
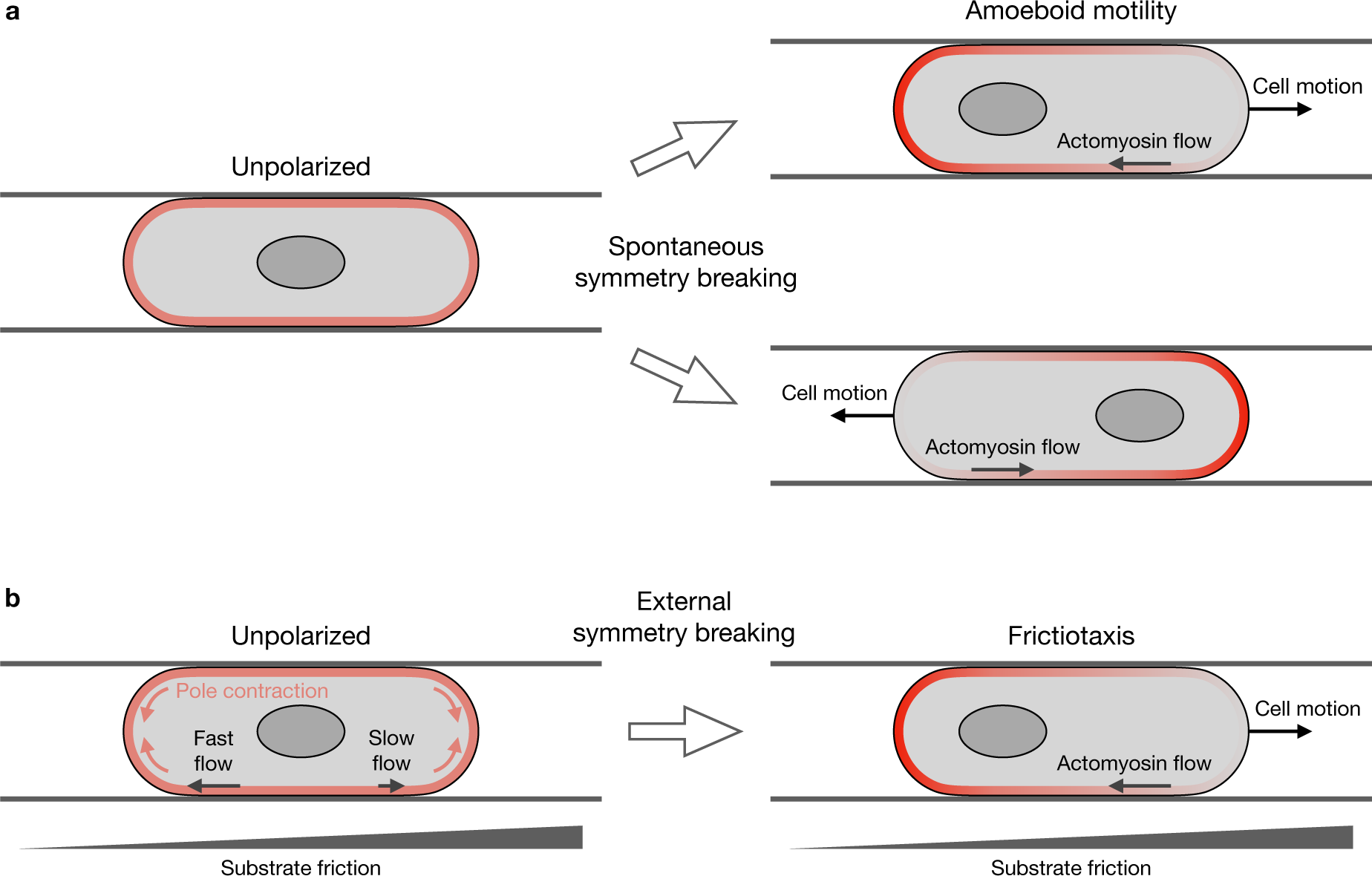
Model of frictiotaxis. **a**, In the absence of external gradients, unpolarised cells break symmetry spontaneously and display amoeboid motility either left or right. **b**, On a friction gradient, contractile stresses from the poles produce faster cortical flows on the lower-friction side. These flows break the symmetry and accumulate actomyosin towards the low-friction side, which drives frictiotaxis, that is directed cell migration towards higher friction.

To benchmark this model, defined by Eqs. S13-S16 in the Supplementary Note, we solved it numerically and obtained the steady-state actomyosin concentration profile (Methods). This profile fits the experimentally measured profile of myosin intensity (Extended Data Fig. 6-7, Table I, see Methods), which shows that the model captures the myosin-driven amoeboid migration in our experiments.

**Table I.** Fit parameter values. Values of the scaling factor *I*_0_ and the pressure coefficient 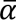 obtained from the fits shown in Extended Data Figs. 6-7, which correspond to 7 cells migrating on uniform friction and 7 cells migrating on a friction gradient, respectively.

| Cell # | Uniform friction |  | Friction gradient |  |
| --- | --- | --- | --- | --- |
| | $I_0/10$ | $\bar{\alpha}/10^2$ | $I_0/10$ | $\bar{\alpha}/10^2$ |
| 1 | $12.0 \pm 0.5$ | $2.6 \pm 0.5$ | $6.3 \pm 0.2$ | $0.5 \pm 0.2$ |
| 2 | $9.5 \pm 0.4$ | $3.3 \pm 0.8$ | $4.7 \pm 0.2$ | $1.3 \pm 0.7$ |
| 3 | $8.9 \pm 0.4$ | $2.0 \pm 1.0$ | $6.2 \pm 0.3$ | $1.8 \pm 0.7$ |
| 4 | $10.1 \pm 0.5$ | $1.4 \pm 0.6$ | $8.0 \pm 0.7$ | $3.0 \pm 3.0$ |
| 5 | $7.8 \pm 0.4$ | $2.0 \pm 2.0$ | $6.9 \pm 0.4$ | $1.2 \pm 0.6$ |
| 6 | $9.0 \pm 0.5$ | $2.0 \pm 1.0$ | $5.2 \pm 0.1$ | $14.0 \pm 1.0$ |
| 7 | $10.9 \pm 0.4$ | $4.0 \pm 2.0$ | $8.5 \pm 0.3$ | $2.2 \pm 0.5$ |

Within the framework of this model, we next asked what would happen if the cells were exposed to an external friction gradient. We show that, in the model, a friction gradient can break the symmetry and yield migration toward regions of higher friction. Starting from an unpolarised state with uniform gel concentration, contractile stresses from the cell poles generate faster cortex flows in low-friction areas. These flows concentrate the gel towards the low-friction side, which thus becomes the cell rear (Fig. 5b).

To demonstrate this mechanism of symmetry breaking, we considered force balance for the actomyosin gel:

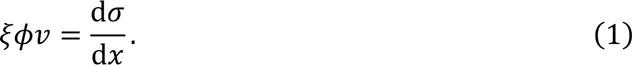

The left-hand side represents gel-substrate friction, which is proportional to the coefficient *ξ*(*x*), the volume fraction ϕ(*x*) of the gel, and its velocity *v*(*x*) with respect to the substrate. The right-hand side represents the forces generated within the gel. These forces arise from gradients of the gel stress σ = *σ_a_* + *σ_v_* − *π*, which includes contributions from active contractility, *σ_a_* = *ζϕ*, gel viscosity *σ_v_* = *ηϕ* d*v*/d*x*, and osmotic pressure *π* = *α*(*ϕ* – *ϕ*_0_)^3^ − γ d^2^*ϕ*/d*x*^224^. Here, *ζ*, *η*, *α*, and *γ* are material parameters described in more detail in Reference^24^ and in the Supplementary Note, and *ϕ*_0_ is the equilibrium volume fraction of the gel.

Because the cell poles are not subject to substrate friction, they undergo the contractile instability earlier and faster than the rest of the gel. The resulting pole contraction pulls in gel from the central region of the cell (Fig. 5b). In the central region, the gel starts in the unpolarised state with uniform gel concentration *ϕ*(*x*) = *ϕ*_0_, for which the force balance Eq. (1) reduces to

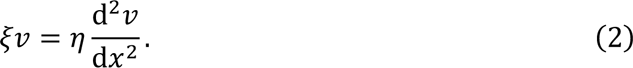

To capture the mechanism of symmetry breaking, we solve this force balance close to the two ends of the central region, which are subject to a stress *σ*_pole_ arising from pole contraction and have different friction coefficients *ξ*___ and *ξ*_+_, with *ξ*_−_ < *ξ*_+_. Thus, we obtain the velocities of the two ends, *v*_−_ and *v*, (Supplementary Note):

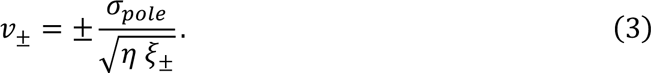

As illustrated in Fig. 5b, this result shows that the gel flows faster towards the pole in the low-friction region, which leads to gel accumulation towards that side and subsequent cell migration towards high-friction regions.

Overall, our theory predicts a mode of cell migration guided by friction gradients (Fig. 5b), which we call frictiotaxis. These results suggest that cells without strong focal adhesions perform durotaxis by exploiting gradients in friction rather than stiffness variations.

## Substrate friction and stiffness are correlated

To test this idea in experiments, we first probed the relationship between stiffness and friction by performing lateral force microscopy (LFM) on the agarose microchannels across the stiffness range in which we observed durotaxis. LFM measures the torsional deformation of the micro-mechanical cantilever of an atomic force microscope during contact mode, enabling the measurement of frictional properties. We found that, in accordance with our hypothesis, stiffer substrates exhibited higher friction forces (Fig. 6a).

**Figure 6.**
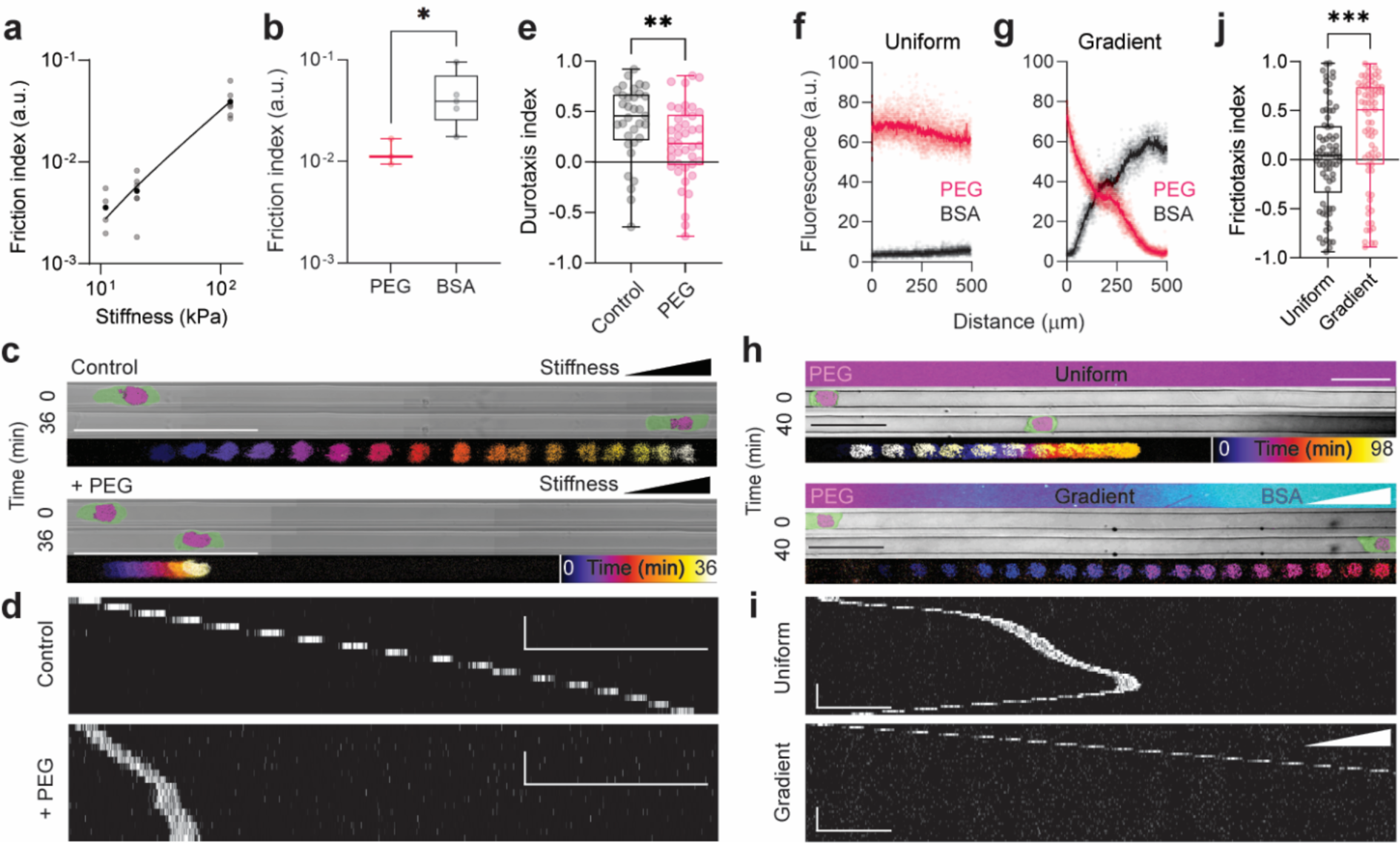
Friction-driven directed migration. **a**, Friction index versus stiffness in the gels; *n* = 4 (*in vitro*, at 11 kPa); *n* = 6 (*in vitro*, other stiffness values). **b**, Friction of PEG- or BSA-treated glass; *n* = 3 (PEG); *n* = 5 (BSA); two-tailed Mann-Whitney test; \**P*≤0.05 (exact: *P*=0.0357). **c**, Pictures of Walker cells with a nuclear label and pseudocoloured membrane after t=0 min and t=36 min in stiffness-gradient microchannels without (Control, upper panel) or with PLL-PEG coating (+ PEG, lower panel). Temporal colour-coded projected tracks of the nucleus are shown in the bottom of each of the panels. Scale bars, 100 µm. **d**, Kymographs corresponding to the cells shown in **c**. Scale bars, 100 µm (horizontal) and 10 min (vertical). **e**, Durotaxis quantification; *n* = 33 cells; two-tailed Mann-Whitney test; \*\**P*≤0.01 (exact: *P* = 0.0097). **f**-**g**, Quantification of micropatterned PEG and BSA in microchannels with uniform and graded friction; *n* = 10 (PEG); *n* = 9 (BSA). Pale dots represent raw data; dark dots represent mean. **h**, Images of PLL-g-PEG/FITC and BSA-Alexa647 micropatterning (top panels), images of cells with a nuclear marker and pseudocoloured membrane at t=0 min and t=40 min (middle panels), and temporal colour-coded projected tracks (bottom panels). Scale bars, 50 µm. **i**, Kymographs corresponding to the cells shown in **h**. Cells were tracked via the nuclear marker. Scale bars, 50 µm (horizontal) and 10 min (vertical). **j**, Quantification of frictiotaxis index from *n* = 74 cells each; two-tailed Mann-Whitney test; \*\*\**P*≤0.001 (exact: *P* = 0.0010).

## Experimental evidence of frictiotaxis

If amoeboid durotaxis is based on friction, it should be impaired if the channel walls are passivated with polyethylene glycol (PEG), which has been shown to provide low-friction substrates^18,26^ (Fig. 6b). Additionally, cells should be capable of directed migration when stiffness is uniform, but friction is graded.

To test the first prediction, we employed a previously described method in which agarose is coupled to proteins after the surface is activated with cyanogen bromide, enabling functional interactions between the cell and the coated surface^27,28^. As a proof of principle, cells adhered strongly to fibronectin-coated agarose gels (Extended Data Fig. 8a). We thus coated agarose microchannels exhibiting a stiffness gradient with PLL-g-PEG (Extended Data Fig. 8b,c). Cells migrating through such channels had impaired durotaxis, compared to controls (Fig. 6c-e). Thus, durotaxis is impaired by a reduced friction despite the presence of a stiffness gradient.

To address whether gradients in friction are sufficient to guide cell migration in the absence of adhesions, we generated gradients in friction by micropatterning gradients of bovine serum albumin (BSA), which provides high friction, and PEG, which provides low friction^18^, onto glass and within PDMS microchannels (Fig. 6f,h and Extended Data Fig. 8d,e), such that regions without BSA contained PEG. By performing LFM, we validated that BSA-coated regions provided higher friction than PEG-coated regions (Fig. 6b), which supports previous evidence^18^. We observed cells moving toward areas of higher friction when they migrated in these channels, as opposed to randomly directed motion when friction was uniform (Fig. 6h-j and Supplementary Video 7). Motion guided by friction gradients was previously conceptualized^29^ and demonstrated in magnetic colloidal particles^30^. Here, our experimental data confirm that cells perform frictiotaxis, and they suggest that this mode of directed migration provides the mechanism for adhesion-independent durotaxis.

## Discussion

Altogether, our study contributes to the growing field questioning how the physical environment modulates adhesion-independent migration^22,31,32^. Our results reveal that strong and specific adhesions are redundant for durotaxis thanks to non-specific friction forces, which enable preferential movement toward stiffer substrates because stiffness and friction are correlated. In practice, different tissues may exhibit different stiffness-friction relationships, thus affecting cell response. Although cells performing amoeboid migration might have weak adhesions, the mechanism of directed motion that we reveal is entirely based on friction. Accordingly, our model includes no adhesion, i.e., no resistance to normal pulling forces, but just friction that opposes cell-substrate sliding.

Modulating only one physical variable, in this case stiffness, to investigate cell behaviour is a major challenge. Despite establishing the dominant role of friction, we cannot rule out that differential substrate deformation contributes to adhesion-independent durotaxis. In this study, we used microchannels that were larger than the nucleus, because amoeboid cells use the nucleus as a mechanical gauge in path-making decisions^33^. We also chose to use preformed paths, rather than an ‘under agarose assay’, to rule out the role of compression on the cells, since cells under compression must physically deform their surroundings to move, which is more difficult when the substrate is stiffer^31^. Whether cell compression and other physical inputs can act as guidance cues for adhesion-independent cell motility remains an open question. Notably, however, geometric patterns can guide adhesion-independent motility^22,34^.

Finally, whether frictiotaxis and amoeboid durotaxis are physiologically relevant in the complex *in vivo* environment, especially considering that chemotactic signals are extremely potent drivers of directional cell migration, is also unknown. A promising candidate are immune cells, which often need to traverse tissues of high density to reach target sites. Adhesion-independent durotaxis could contribute significantly to this type of directed migration.

---

**Extended Data Fig 1.**
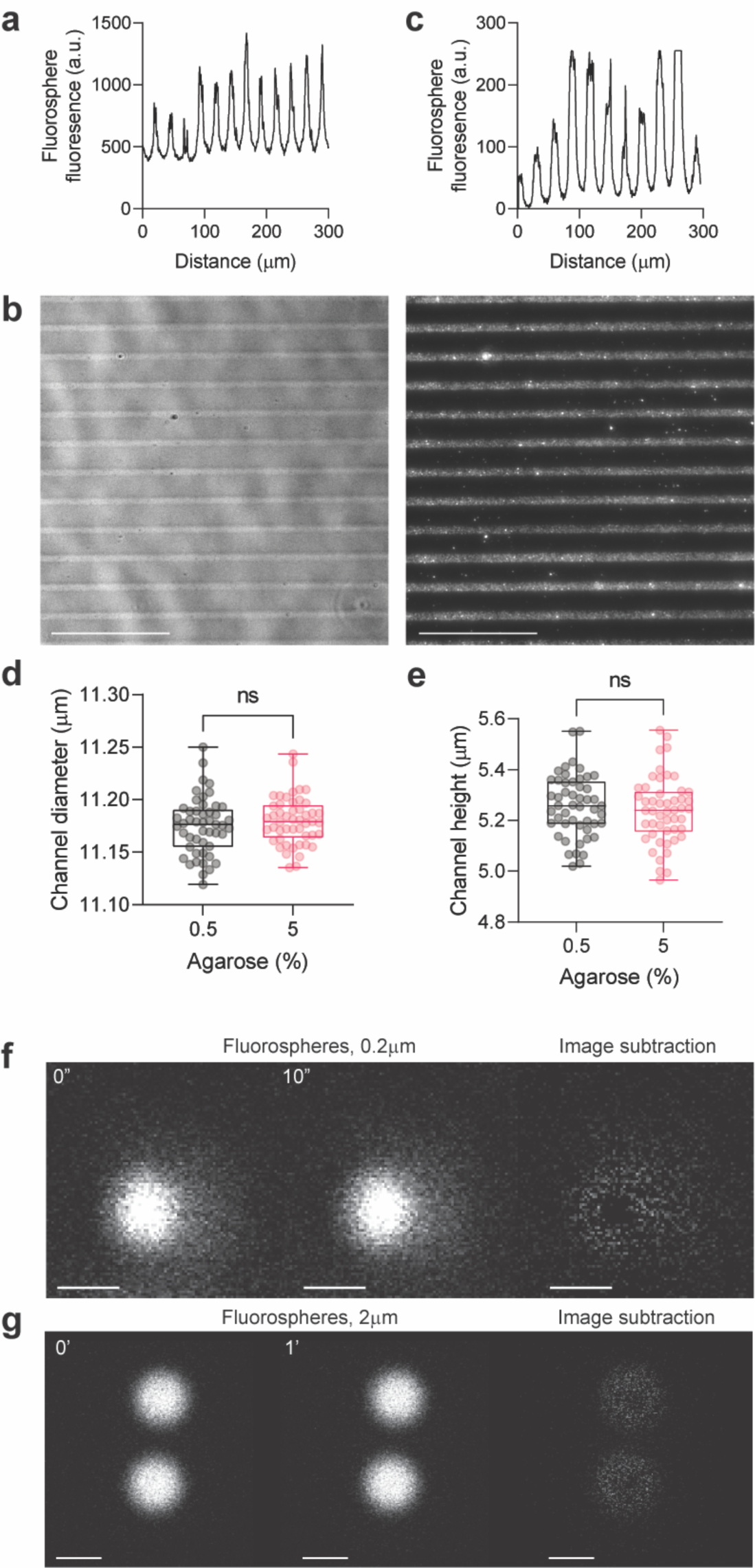
The agarose microchannel assay. **a-c**, 5 µm agarose channels filled with 0.2 µm fluorospheres (**b**) and quantification along the orthogonal axis of fluorosphere-filled 10 µm (**c**) and 5 µm channels (**a**), as in Fig. 1c and Extended Data Fig. 1b, respectively. Scale bar, 100 µm. **d-e**, Channel dimensions quantified. Translucent dots represent individual data points. *N* = 50 channels; unpaired two-tailed t test; ns, *P*>0.05 (exact: *P*=0.3482 in **d**; *P*=0.4649 in **e**). **f-g**, 0.2 µm (**f**) or 2 µm fluorospheres (**g**) in 10 µm agarose microchannels at different time points. The right panel is an image subtraction between the images. Scale bar, 0.5 µm (**f**); 2 µm (**g**).

**Extended Data Fig 2.**
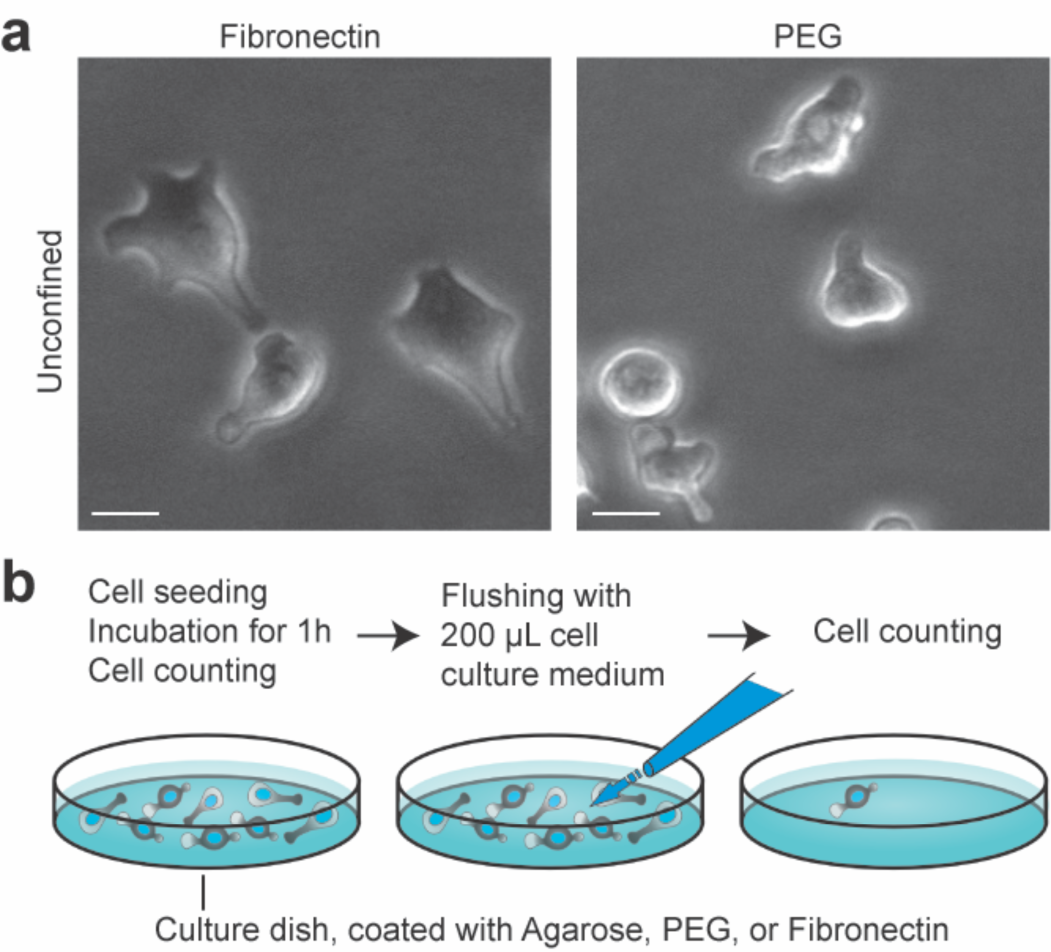
Walker cells lack strong and specific adhesions. **a,** Pictures of cells in adhesive (fibronectin, left) or non-adhesive (PEG, right) conditions. Scale bar, 10 µm. **b**, Illustration depicting the principle and quantification of the flushing experiment.

**Extended Data Fig 3.**
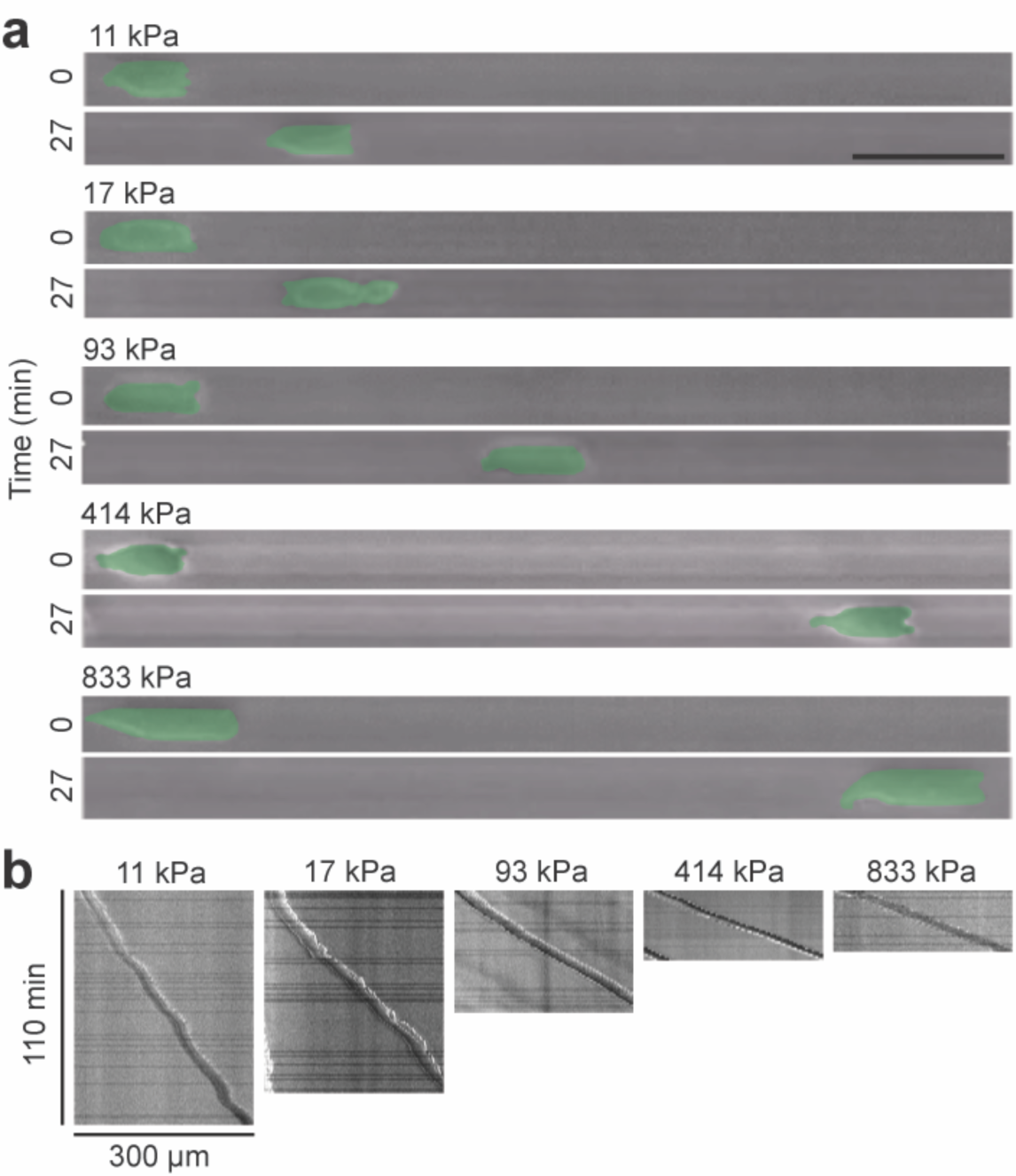
Stiffness-dependent migration speed in microchannels. **a**, Pictures of cells with pseudocoloured membrane at early (t=0 min) and later (t=27 min) time points in microchannels of different stiffness regimes. Scale bar, 50 µm. **b**, Kymographs corresponding to the cells depicted in (**a**). Scale bars equal 300 µm (horizontal) and 110 min (vertical).

**Extended Data Fig 4.**
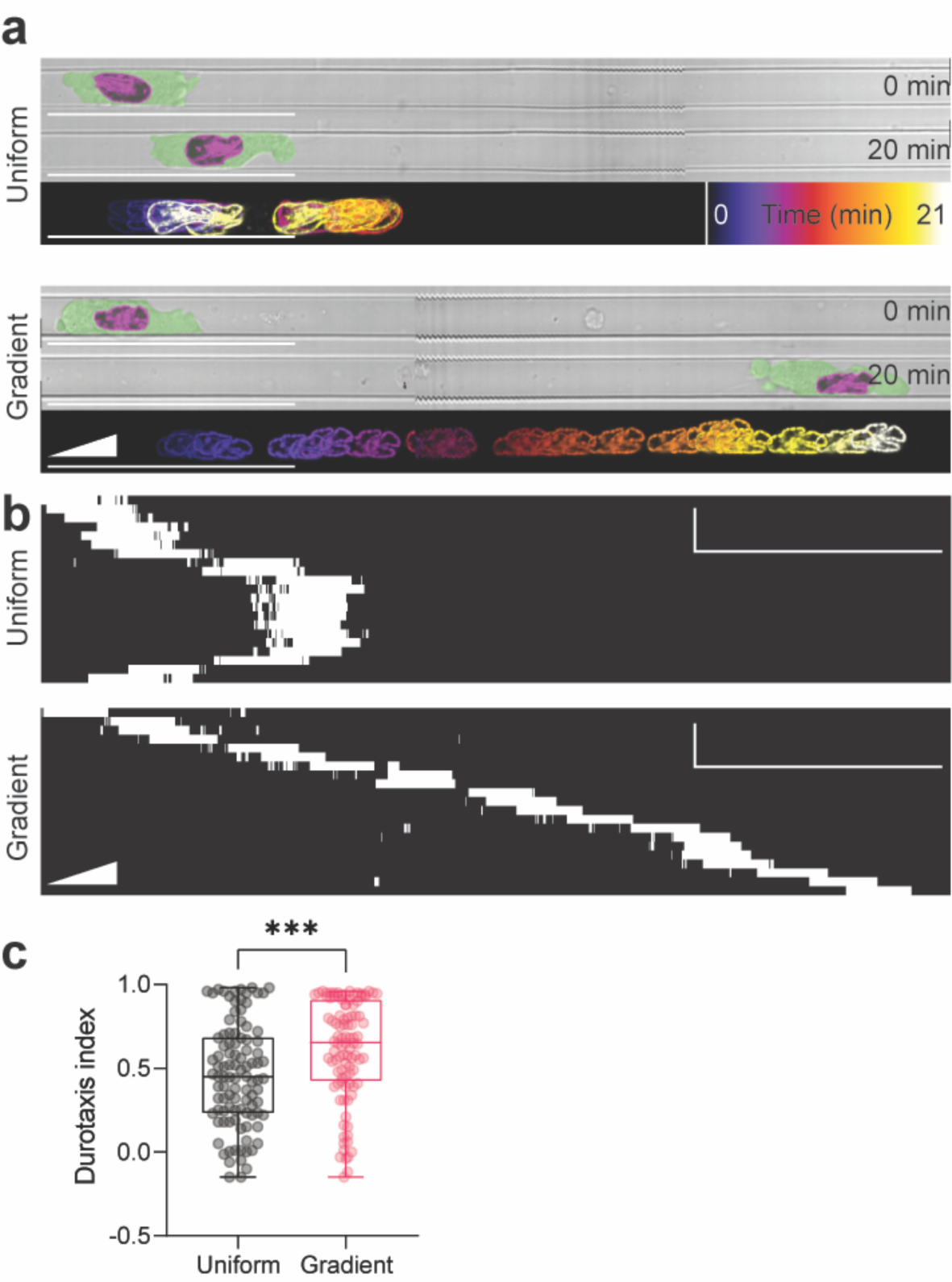
Durotaxis of HL60 cells. **a-b,** Example cells at early and later time points (upper and middle panels, **a**), temporal colour-coded projections (bottom panels, **a**) and kymographs (**b**) in uniform and graded stiffness substrate. Cells were tracked by labelling the nucleus with Hoechst 33342 and the cell membrane was pseudocoloured. Images are stitched as described in the Methods. Scale bar, 50 µm. Vertical scale bar (**b**), 5 min. **c,** Durotaxis quantification. *n* = 100 cells; two-tailed Mann-Whitney test; \*\*\**P*≤0.001 (exact: *P*=0.0006).

**Extended Data Fig 5.**
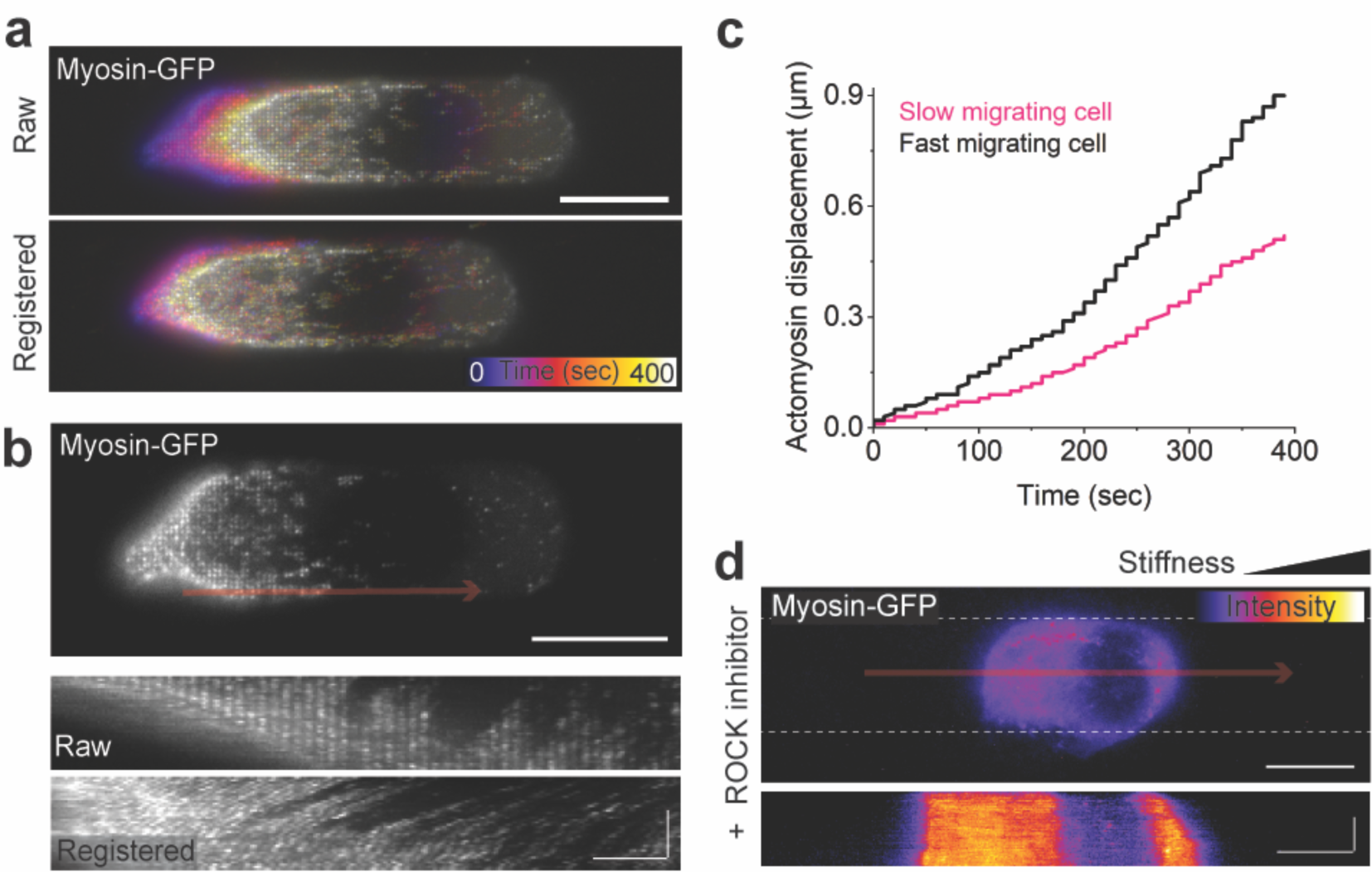
Visualisation of retrograde actomyosin flow. **a**, Temporal colour-coded projection of a Myosin-GFP-expressing Walker cell before (raw_top panel) and after image registration (registered_bottom panel). Scale bar equals 10 µm. **b**, Kymographs of actomyosin flow, corresponding to the area along the red arrow in the upper panel, before (raw) and after image registration (registered). Scale bars, 10 µm in the upper panel; 5 µm (horizontal) and 200 s (vertical) in kymographs of lower panels. **c**, Examples of actomyosin displacement over time of a slow (1.47 µm/min) versus faster (2.16 µm/min) migrating cell in a microchannel with stiffness gradient. **d**, Colour-coded still image (upper panel) and Kymograph (lower panel, corresponding to the red arrow in the upper panel) of a Walker cell expressing Myosin-GFP, in a microchannel with stiffness gradient, and treated with 30 µM ROCK inhibitor (Y-27632). Dotted lines in the upper panel represent channel boundaries. Scale bars, 10 µm in upper panel; 10 µm (horizontal) and 20 min (vertical) in lower panel.

**Extended Data Fig 6.**
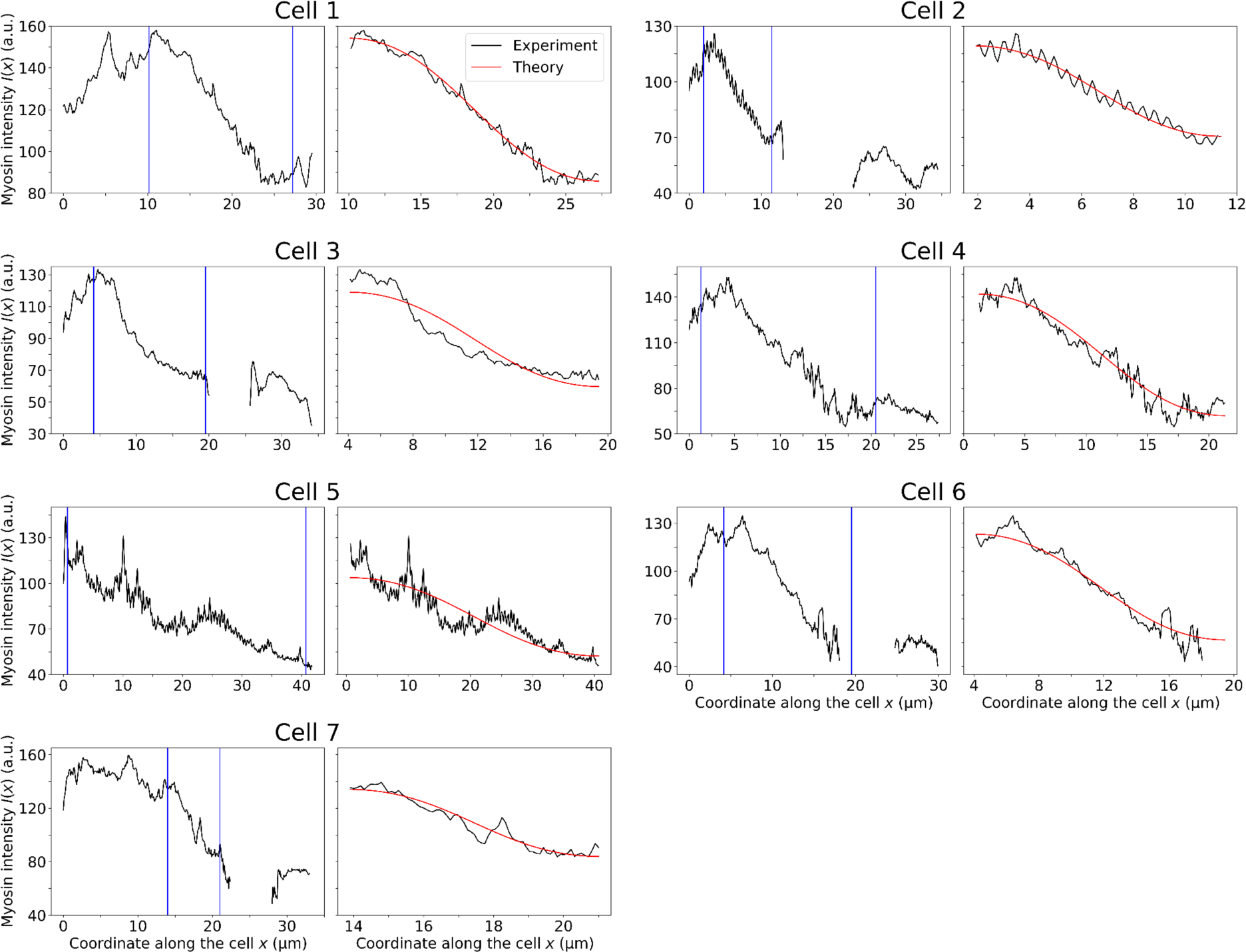
Fits of the myosin intensity profiles for cells migrating on uniform friction. For each cell, the left column shows the myosin intensity profile along the cell at a single time frame. For some cells, there is missing intensity data at certain positions due to the nucleus displacing myosin. Only the data between the vertical blue lines is used for comparison to the steady-state solutions of the model. For each cell, the right column shows the fit of the steady-state solutions of the model (red) to the experimental data (black) selected from the left panels. Error bars are not visible as the errors of the mean of the experimentally measured intensities are typically around 1-5 a.u., which is much less than their absolute values. The fit parameter values are listed in Table I.

**Extended Data Fig 7.**
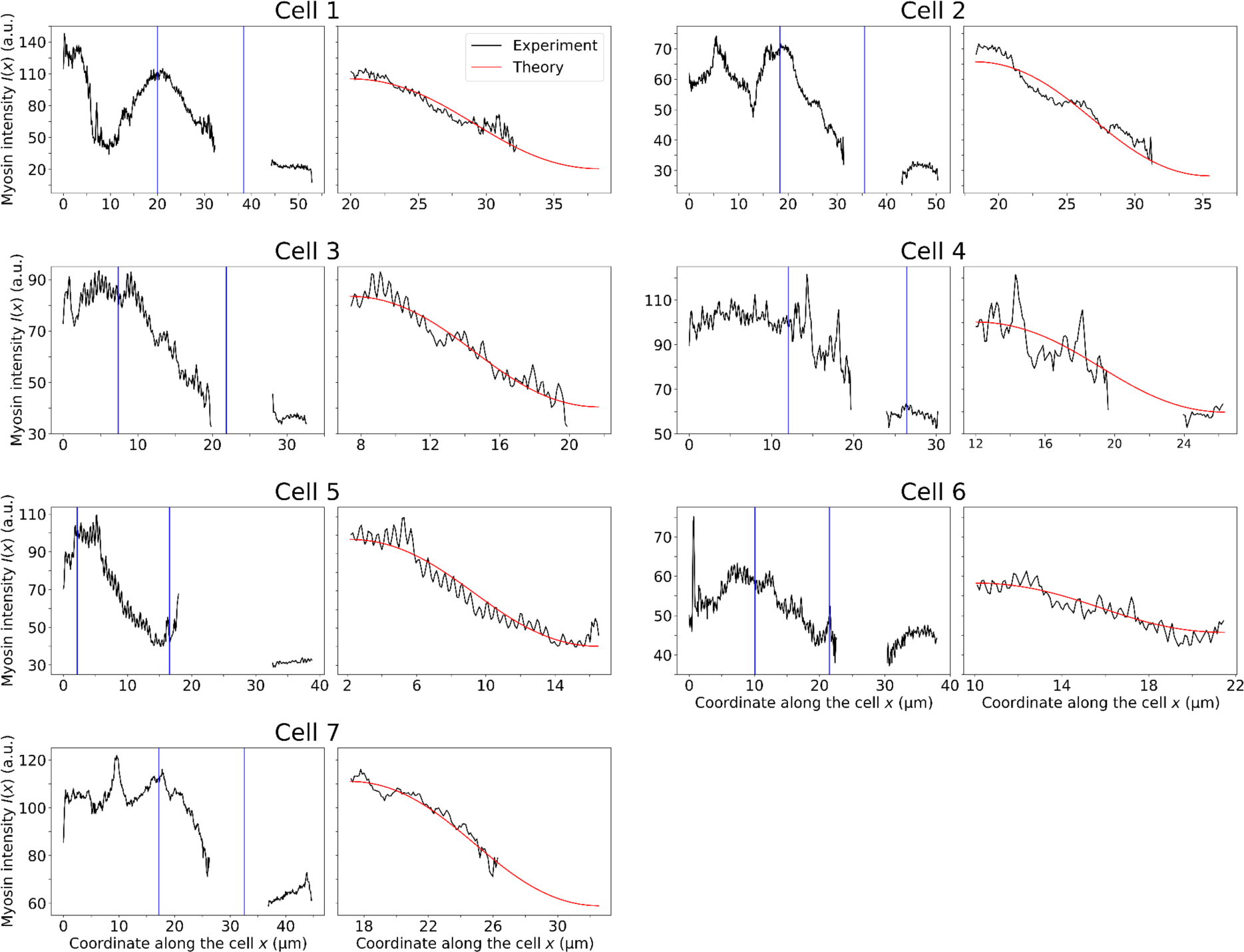
Fits of the myosin intensity profiles for cells migrating on a friction gradient. For each cell, the left column shows the myosin intensity profile along the cell at a single time frame. For some cells, there is missing intensity data at certain positions due to the nucleus displacing myosin. Only the data between the vertical blue lines is used for comparison to the steady-state solutions of the model. For each cell, the right column shows the fit of the steady-state solutions of the model (red) to the experimental data (black) selected from the left panels. Error bars are not visible as the errors of the mean of the experimentally measured intensities are typically around 1-5 a.u., which is much less than their absolute values. The fit parameter values are listed in Table I.

**Extended Data Fig. 8.**
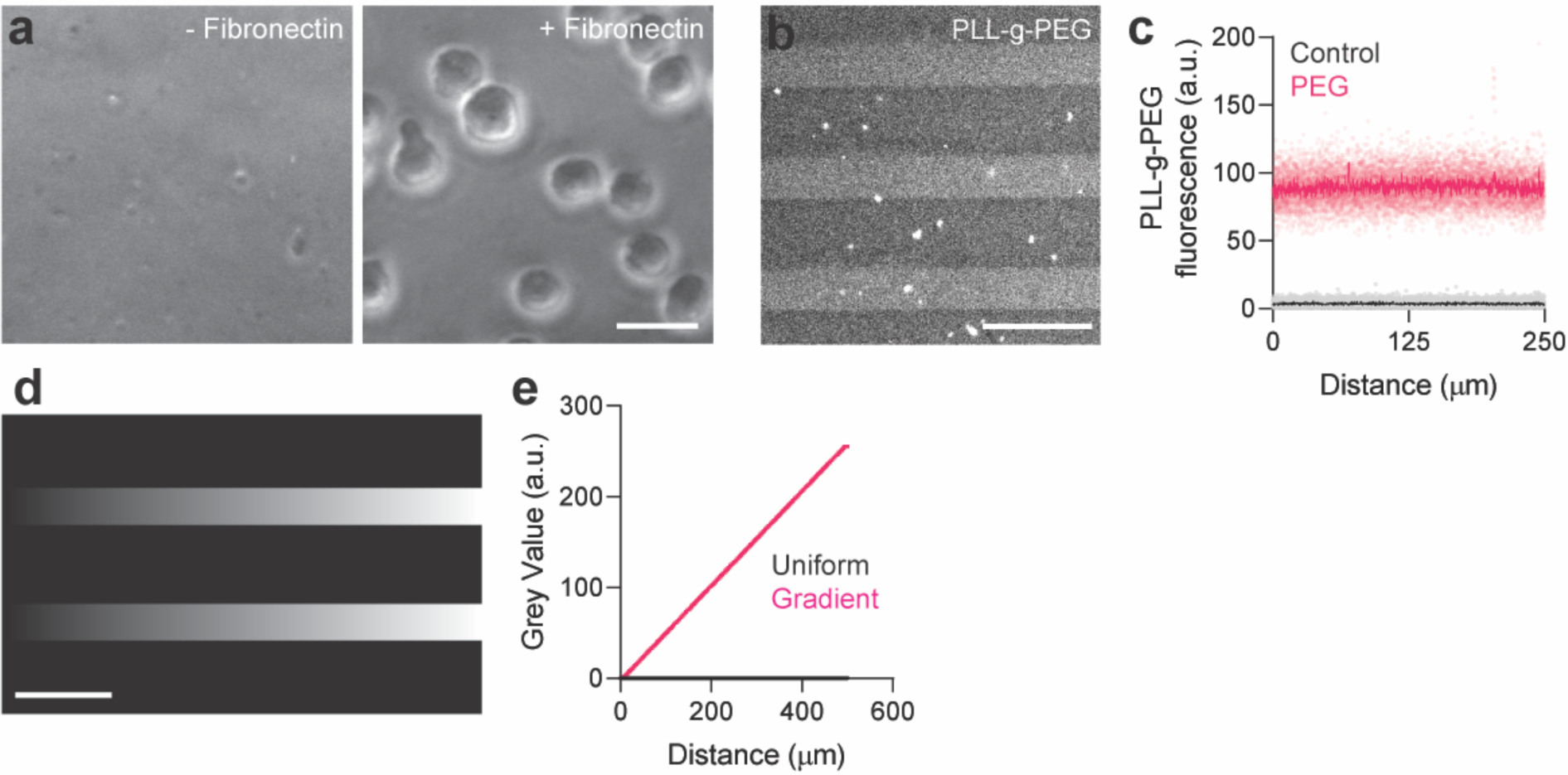
Agarose activation, PEG coating and photopatterning gradients for frictiotaxis experiments. **a,** Cells adhere to a fibronectin-coated agarose gel. Medium was aspirated and flushed several times to remove any cells in suspension. Scale bar, 20 µm. **b-c**, PLL-g-PEG/FITC-coated or uncoated (control) agarose microchannels (**b**) and quantification of fluorescence (**c**); *n* = 10; pale colours represent raw data; dark colours represent mean. Scale bar, 20 µm. **d**, The designed photopattern which exhibits alternating control and gradient masks. Scale bar, 100 µm. **e**, Quantification of (**d)**.

**Supplementary Video 1. Agarose microchannel assay.**

Walker cells migrating within 10 µm width microchannels within a 1% agarose substrate. Movie length, 481 min. Scale bar, 100 µm.

**Supplementary Video 2. Adhesion-independent durotaxis.**

Walker cells undergo durotaxis without strong or specific adhesions to the substrate. Movie length, 40 min. Scale bar, 50 µm.

**Supplementary Video 3. Stiffness-dependent migration speed.**

Migration speed of Walker cells increases with higher substrate stiffness. Movie length, 110 min.

**Supplementary Video 4. Myosin rear accumulation in gradient and uniform stiffness.**

In stiffness gradient and uniform stiffness microchannels, Myosin-GFP accumulates at the cell rear and accumulation switches to the opposite pole when cells repolarise.

**Supplementary Video 5. Retrograde Myosin-GFP flow in raw and registered sequences.**

Retrograde Myosin-GFP flow is observed in raw and registered timelapse sequences.

**Supplementary Video 6. Myosin inhibition abrogates migration in microchannels.**

Upper panel: DMSO control. Lower panel: 30 µM Y-27632. Note that both cells in the lower panel do not form blebs and do not migrate.

**Supplementary Video 7. Frictiotaxis.**

Walker cells undergo frictiotaxis on PEG-BSA gradients. Movie length, 98 min. Scale bar, 50 µm.

## Materials and Methods

### Cell culture

Walker 256 carcinosarcoma cells (RRID:CVCL_4984), Walker myosin light chain-GFP cells, and Walker Lifeact-GFP cells were a gift from E. Paluch (University of Cambridge, UK). Walker cells and HL60 cells (RRID:CVCL_0002) were grown in T-25 suspension cell culture flasks (Sarstedt, 83.3910.502) in RPMI 1640 media containing L-glutamine (Gibco, 11875101) supplemented with 10% heat-inactivated FCS and 1% penicillin-streptomycin (Gibco, 10378016) at 37°C and 5% CO2. The culture media for the HL60 cells additionally contained 40 mM Hepes for buffering. Differentiation of HL60 cells was achieved by incubating cells in propagation media plus 1.3% DMSO for 5 days. All working media was passed through 0.2 µm Sartorius Minisart filters (VWR, 611-0691) prior to use. For experiments, cells were transferred to media identical to their culture media except lacking FCS. For myosin inhibition experiments, the imaging medium was additionally supplemented with 30 µM of the ROCK inhibitor Y-27632 (Selleckchem, S6390), and an equivalent volume of DMSO was added to the control condition. Cells were allowed to equilibrate in the medium for 30 min before the start of the experiment.

Where appropriate, cells were labelled with Hoechst 33342 (Leica, H3570) at 1×10^−5^ mg/mL, or BioTracker 490 Green Cytoplasmic Membrane Dye (Merck, SCT106) at 5 µL per 1 mL cell suspension. In both cases, a concentrated cell solution was incubated with the marker for 30 min at 37°C before being washed out. Immunostaining against phospho-paxillin Tyr 118 (ThermoFisher, 44-722G) was performed by fixing in 4% paraformaldehyde for 10 min at 37°C followed by 0.1% Triton X-100 permeabilization for 15 min at RT, blocking in 2% BSA/PBS for 1 hour at RT, 1:1000 primary antibody overnight at 4°C and a AlexaFluor goat anti-rabbit secondary antibody (ThermoFisher, A-11008) at 1:500 for 4 hours at RT with PBS washes in between antibody steps.

### Image processing, data analysis and statistics

Images and videos were processed using Fiji. Cells were tracked using the Manual Tracking plugin, and output data processed in the Chemotaxis and Migration Tool (Ibidi, Version 2.0). In this manuscript, we have named Forward Migration Index as the Durotaxis Index, where the axis of interest is along the stiffness gradient and where positive values indicate straightness toward stiff substrate, and negative values indicate straightness toward soft substrate.

To achieve high resolution imaging of cells across large regions, images were stitched for analysis and presentation. Correction for uneven illumination was performed where appropriate. Cells in contact with other cells were not included in the analysis, and tracks were cut short where cell-cell contacts were made, to ensure only single cell analysis was performed.

To quantify retrograde actomyosin flow independently from cell displacement, linear stack alignment with SIFT was carried out in Fiji, and displacement was tracked manually in kymographs, using the segmented line tool.

Normality in the spread of data for each experiment was tested using the Kolmogorov-Smirnov, d’Agostino-Pearson, and Shapiro-Wilk tests in Prism9 (GraphPad9). Significances for datasets displaying normal distributions were calculated in Prism9 with paired or unpaired two-tailed Student’s *t*-tests or Mann-Whitney U tests where appropriate. No predetermination of sample sizes were done. Cells were allocated into experimental groups randomly. Authors were not blinded because cells were selected prior to analysis. Criteria for selection was survival and not interacting with other cells. All experiments were replicated three times (biological replicates) unless otherwise indicated.

### Nanoindentation

Stiffness measurements were performed using nanoindentation (Chiaro, Optics11Life, Piuma V2 v3.4.3) as previously described^3^. Cantilevers were customized by Optics11 Life. Probes had a spherical glass tip with a radius of ∼ 10 µm mounted onto an individually calibrated cantilever with a spring constant of ∼ 0.25 N m^−1^. Deformation of the cantilever after contact with the sample was measured by tracking the phase-shift in light, reflected from the back of the cantilever. Samples were indented to a depth of 0.5 μm with a velocity of 2.5 μm s^−1^. The tip was held at this indentation depth for 1 s and then retracted over 1 s. The Young’s moduli were calculated automatically by the software by fitting the force versus indentation curve to the linear Hertzian contact equation model^35^. The effective Young’s modulus (*E*), referred to in this manuscript as stiffness, is derived from the fit of the loading force-displacement curve (*F*(*h*)), the indenter tip radius (*R*), and the indentation depth (*h*), according to the following formula, for which a Poisson’s ratio (*v*) of 0.5 was assumed, and was calculated automatically by the software (Chiaro, Optics11Life, Piuma V2 v3.4.3).

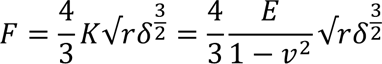

### Atomic force microscopy

10 µm polystyrene beads were glued onto tipless MLCT B cantilevers (nominal spring constant 0.02 N/m) and friction measurements were performed on the Nanowizard 4 AFM system (Bruker Nano GmbH, JPK BioAFM) using Contact Mode. Lateral force was recorded by sweeping the cantilever over a line of 10 µm length at a speed of 2.5 µm/s. We measured the force at 5 kHz over a time frame of approximately 60 seconds to record several oscillations. For each gel forces of 0.25, 0.5, 0.75, 1, 1.25, 1.5 and 1.75 nN was applied and repeated at each force 5 times.

### Micropatterning

PDMS or glass was plasma treated (Diener electronic) for 2 min and immediately coated with 100 µg/ml PLL(20)-g[3.5]-PEG(2) (Susos) or PLL(20)-g[3.5]-PEG(2)/FITC (Susos), dissolved in PBS, for 30 min at room temperature. Surfaces were washed with 100 mM Hepes pH 8-8.5 for 1h at room temperature and then incubated with 100 mg/ml mPEG-SVA (Laysan Bio, MPEG-SVA-5000). Surfaces were washed in 1x PBS, followed by water, and then air dried. The photoactivatable reagent, PLPP gel (Alvéole) was added at a ratio of 3:17 with 70% ethanol, at 1 µl PLPP gel/cm^2^, and allowed to dry completely whilst protected from light.

PRIMO (Alvéole) was calibrated with a 20x objective on a Nikon Ti inverted microscope using Leonardo software (Alvéole). A greyscale pattern was used as shown in Extended Data Fig. 8d. A dose of 30 mJ/mm^2^ was used and laser power adjusted such that the patterning time for each element was 4 s.

After patterning, the surface was washed several times with water and re-hydrated with PBS for 5 min. The sample was then incubated with 50 µg/ml AlexaFluor647-conjugated BSA (Thermo Fisher, A34785) overnight at 4°C prior to washing with PBS. Subsequent stages of PDMS bonding to glass and imaging were performed as described in the Methods sections, Soft lithography, and Imaging.

### Photolithography

3-inch silicon wafers (Silicon wafer test grade, N(Phos), WAFER-SILI-0580W25 from PI-KEM Ltd.) were plasma cleaned for 10 min at 100% power in a plasma cleaner (Henniker Plasma HPT 100), and then baked on a hot plate at 200°C for 20 min to remove moisture. After 10 s of cooling, SU-8 2005 (Kayaku Advanced Materials, purchased from A-Gas Electronics Materials) was spin coated on the wafer at 500 rpm with an acceleration of 100 rpm s^−1^ for 30 s. After 2 min rest, the wafer was soft baked at 65°C for 1 min, followed by 95°C for 2 min, and then allowed to cool for 10 s. Writing was performed on a MicroWriter ML3 using a 20x objective. Post-exposure bakes of the wafer were then performed at 65°C for 1 min, 95°C for 2 min and then 65°C for 1 min. After 10 s cooling, the wafer was developed in PGMEA (Sigma-Aldrich 484431-1L) for 1 min with manual agitation, before rinsing in IPA for 10 s, dried with compressed N2 and hard baked for 20 min at 200°C. Dimensions of the wafer were analysed on an optical profiler (Sensofar S Neox).

### Soft lithography

PDMS was made using reagents from a SYLGARD 184 Elastomer Kit (VWR, 634165S). Base elastomer was thoroughly mixed with curing agent at a ratio of a ratio of 10:1, before degassing in a vacuum desiccator (Scienceware, 999320237) with a KNF Laboport N96 diaphragm vacuum pump (Merck, Z675091-1EA). A Si wafer with positive features was placed in a disposable aluminium dish (VWR, 611-1377), covered in PDMS, and baked for 10 min at 110°C. The PDMS was separated from the wafer and trimmed using a blade.

### Agarose microchannels

PDMS was prepared as above, and spin coated onto a Si wafer with negative features using a spin coater (Spin150) at conditions of 400 rpm for 30 s. The wafer was then baked at 110°C for 10 min. The thin PDMS was peeled off from the wafer and cut into smaller pieces using a razor blade, where each individual piece contained between one and four patterned designs. To fabricate agarose microchannels, an individual PDMS piece was placed on a glass slide with an edge aligned with the edge of the slide. A vacuum, and subsequently finger pressure, was used to push trapped air bubbles out from underneath the PDMS. A ∼1 mm thick piece of PDMS was cut to generate a U-shape and placed around the PDMS pattern on the slide. A coverslip was placed on top of the U-shape PDMS and neodymium magnets above the coverslip and below the slide were used to secure the sandwich in place.

UltraPure low melting point agarose (ThermoFisher, 16520050) – in this manuscript referred to as ‘agarose’ – was dissolved in RPMI media at the appropriate concentration, depending on the experiment. The solution was maintained at 70°C while penicillin-streptomycin (Gibco, 10378016) was added to a final concentration of 1%. To fabricate gels of uniform stiffness, the agarose solution was pipetted into the sandwich until full. To fabricate gels of graded stiffness, higher concentration agarose solution (4%) was pipetted into the sandwich mid-way, followed by lower concentration agarose solution (1%) until full. The gradient was modulated by modifying the diffusion rate, which could be controlled by incubating in an oven at different temperatures, angles and by modifying the input solutions. To solidify the agarose gels, the sandwich was transferred to a 4°C fridge. Afterwards, the sandwich was disassembled and the solidified agarose slowly slid away from the underlying PDMS mould, and reversed in orientation such that the structured agarose was face up. The surface was then dried with a nitrogen gun and trimmed to size with a razorblade. The stiffness of agarose gels was measured using a Chiaro nanointender (Methods, nanoindentation).

Holes were punched using a Harris uni-core 3 mm biopsy puncher (VWR, 89022-356), followed by air drying with a nitrogen gun. A glass coverslip was then plasma cleaned and bonded to the agarose microchannel chip, with the channels facing the coverslip. RPMI media was pipetted into the holes and the unit placed in a humid chamber at 37°C and 5% CO2 for 30 min. The wells were then emptied using a p200 pipette and replaced with concentrated cell solution. A 0.5 g weighted glass slide was placed on top of the structure, and the unit placed in the stage of an inverted microscope.

Where appropriate, agarose surfaces were covalently bound to protein by using the CNBr-method that has been reported previously^27,28^. After drying, agarose microchannels were activated with cyanogen bromide: 50 mg/ml in water was mixed in an equal ratio with 0.5 M Na2CO3 in NaOH buffer, pH 11, which contained the protein (fibronectin or PLL-g-PEG). After 30 min, the surface was washed with water and then with the coupling buffer: 0.1 M sodium borate buffer, pH 8.5 for 4 h.

### Measuring the myosin intensity profile

Fluorescent live cell imaging was performed on a ZEISS Elyra 7 microscope equipped with a 40×, NA = 1.2 water-immersion objective and operating in laser WF mode. Time series of single plane images along the cell-agarose interface were acquired in intervals of 10s. To prevent GFP quenching, RPMI medium without phenol red (Thermo Fisher, 11835030) was used in all experiments. Images were exported as TIF series for further analysis. The experimental snapshots showed an intense myosin-GFP signal across the entire cell area except for the nuclear region. We cropped individual snapshots by hand using Fiji to exclude the background and nuclear region. We averaged the pixel intensities along the axis perpendicular to the direction of motion to obtain a one-dimensional intensity profile *I*(*x*) along the cell.

### Numerical solution of the active gel model for amoeboid migration

To reduce the number of free parameters, we simplified the model. Firstly, we assumed that the friction coefficient was constant. The equations in the model can be made independent of the equilibrium volume fraction *ϕ*_0_. This can be seen by rescaling the gel volume fraction *ϕ* → *ϕ*_0_*ϕ*, the pressure coefficient 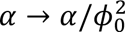 and the drag coefficient *ξ* → *ξ*_f_*ϕ*_0_. We therefore set *ϕ*_0_ = 1 without loss of generality. Finally, we made Eqs. S13-16 in the Supplementary Note dimensionless as in Reference^24^. In dimensionless form, the model contains 4 dimensionless parameters: the active stress coefficient 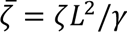, the depolymerization rate 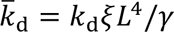, the pressure coefficient 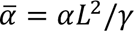, and the drag coefficient 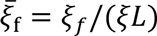.

We solved the dimensionless equations using the Method of Lines^36–38^ to predict the time variation of the steady-state gel volume fraction profile *ϕ*_ss_(*x*, *t*) and the cell velocity *V*(*t*) for migrating cells. We used centered finite differences to approximate the spatial derivatives. Fictitious points were added outside the domain to define the spatial derivatives at the boundaries. We used an explicit Euler step for the equations involving a time derivative and we solved the remaining equations algebraically. We used 25-35 discretization points and a time step of *Δt* = 10^−7^. As initial conditions, we took the homogeneous state *ϕ*(*x*) = *ϕ*_0_ plus a small Gaussian-noise perturbation to each variable. The algorithm converges to provide a steady-state gel volume fraction *ϕ*_ss_(*x*) and the corresponding cell velocity *V*_ss_.

### Fitting the model to the myosin intensity profiles

We take the myosin intensity profile *I*(*x*) from the experimental images as a proxy for the concentration of the actomyosin cortex, given by its steady-state volume fraction *ϕ*_ss_(*x*) in the active gel model. However, we first had to select the region of the myosin intensity profile that can be captured by the active gel model, given its assumptions. The model captures cortical flow from low to high cortex concentrations and it assumes that the concentration gradient vanishes at the rear and front of the cell. In contrast, experimental myosin intensity profiles typically show extended intensity plateaus at each end of the cell, which are connected by an intermediate intensity decay. To match the boundary conditions of the model, we cropped the myosin intensity profiles to include only the intermediate decay region and a small connected region of each plateau. In some cases, the myosin intensity at the start of the plateau is obscured by the nucleus, so we roughly inferred where to crop the profile by extrapolation. Also, cells 1 and 2 in Extended Data Fig. 6 have pronounced uropods, which are myosin-rich tail structures that manifest as a second peak in the myosin intensity profile at the rear side. We did not include the region of the uropod in the comparison to the model. We indicate which region of each myosin intensity profile is selected by means of vertical blue lines in Extended Data Figs. 6-7 (left-hand panels for each cell).

We also had to scale the dimensions of the gel volume fraction *ϕ*(*x*) so that it could be directly compared to the myosin intensity profile *I*(*x*) in the selected region. The coordinate along the cell was scaled linearly as *x* → *xL* to match the length *L* of the selected region. Respectively, to fit it to the myosin intensity profile *I*(*x*), we scaled the gel volume fraction as *I*(*x*) = *I*_0_*ϕ*_ss_(*x*), where *I*_0_is an additional free parameter in the fit.

Before performing the fits, we reduced the number of free parameters in the model. Firstly, we fixed some of them: 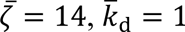, and 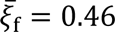. These parameter values ensure that the homogeneous state is only unstable to perturbations with a wavelength given by the cell length. As a result, in this parameter regime, the model has steady-state gel volume fraction solutions *ϕ*_ss_(*x*) that decrease monotonically from the rear to the front of the cell. In addition, each solution has an average value of gel volume fraction that is close to the equilibrium gel volume fraction *ϕ*_0_ = 1. Therefore, for each snapshot, we calculated *I*_0_by measuring the average myosin intensity in the selected region. In some cases, there was missing intensity data in part of the selected region, due to the nucleus obscuring the myosin signal. Therefore, we removed the intensity data at the opposite side of the selected region in calculating *I*_0_, which would otherwise skew the result.

By varying 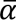, we performed a least-squares fit of the scaled steady-state volume fraction to the myosin intensity profile in the selected region for each cell at a single time frame. We show the fits in Extended Data Figs. 6-7 (right hand panels for each cell), and the corresponding parameter values in Table I.

## Acknowledgements

We thank E. Paluch for gifting us Walker 256 carcinosarcoma cells, L. Alvizi for advice with tissue culture and J. Hartmann for assistance with stitching. We thank P. Saez for advice about PDMS and microchannels. We thank Alveole and Cairn for providing access to the PRIMO, as well as P. March for access to the PRIMO at the University of Manchester. P.H. and R.A. thank A. Callan-Jones, P. Haas, and J. Neipel for discussions on the model and M. Bovyn for discussions on the fits.

## Funding

Work in the laboratory of R.M. is supported by grants from the Medical Research Council (MR/S007792/1), Biotechnology and Biological Services Research Council (M008517; BB/T013044) and Wellcome Trust (102489/Z/13/Z). K.W. is supported by the Deutsche Forschungsgemeinschaft (DFG) via the Walter Benjamin Fellowship (WE 7331/1-1).

## Author contributions

A.S. and K.W. conceived the project, performed the experiments, and analysed the experimental data. R.A. and P.H. developed the physical model and derived the explanation of frictiotaxis. P.H. solved the model numerically and fitted it to the experimental data. N.S. contributed to conceptualisation of the model and performed AFM with R.T. with assistance from G.C. G.C. provided conceptual and technical advice, especially with AFM and tissue culture. A.S and C.D. generated the silicon wafers in A.I.’s laboratory. A.S., K.W., R.M. and R.A. wrote and edited the manuscript. All authors commented on the manuscript.

## Competing interests

The authors declare that they have no competing interests.

## Data and materials availability

All data are available in the main text or the supplementary materials.

## Notes

### Competing Interest Statement

The authors have declared no competing interest.

### Summary of Updates

Cells move directionally along gradients of substrate stiffness, a process called durotaxis. The current consensus is that durotaxis relies on cell substrate focal adhesions to sense stiffness and transmit forces that drive directed motion. Therefore, focal adhesion independent durotaxis is thought to be impossible. Here, we show that confined cells can perform durotaxis despite lacking strong or specific adhesions. This durotactic migration depends on asymmetric myosin distribution and actomyosin retrograde flow. We show that the mechanism of this adhesion-independent durotaxis is that stiffer substrates offer higher friction. We propose a physical model that predicts that nonadherent cells polarise and migrate towards regions of higher friction a process that we call frictiotaxis. We demonstrate frictiotaxis in experiments by showing that cells migrate up a friction gradient even when stiffness is uniform. Our results broaden the potential of durotaxis to guide any cell that contacts a substrate and reveal a new mode of directed migration based on friction, with implications for immune and cancer cells, which commonly move with nonspecific interactions.

