## Supplementary note for "Frictiotaxis underlies adhesion-independent durotaxis"

In this Supplementary Note, we provide further details of our theory for frictiotaxis. For reference, we first review the model of amoeboid cell motility introduced in Ref.[1]. We then use it to argue how cells might polarize in response to a friction gradient.

### A. Active gel model of amoeboid cell motility

The model proposed in Ref.[1] explains amoeboid motility under confinement as a result of a contractile instability of the actomyosin network. As a consequence of this instability, actomyosin becomes enriched at one side of the cell, which becomes the cell rear. The higher concentration drives retrograde actomyosin flow which, through friction with the confining walls, produces forward cell motion (Fig. 5a).

#### 1. *Gel mass balance*

The model treats the actomyosin network as an active gel with volume fraction $\phi(x,t)$. We consider spatial dependence and motion only along the $x$ axis, and the cell spans from $x = 0$ to $x = L$. The model is based on equations for the balance of mass and forces in the gel. Mass balance is given by

$$\partial_t\phi - V\partial_x\phi + \partial_x(\phi v) = a v_{\rm p} - k_{\rm d}\phi = -k_{\rm d}(\phi - \phi_0). \quad \text{(S1)}$$

The second term on the left-hand side captures the gel motion at the cell velocity $V$, corresponding to a travelling gel concentration profile of the form $\phi(x-Vt)$. The third term corresponds to the advection of gel by its velocity $v$. The right-hand side accounts for gel polymerization at a speed $v_{\rm p}$ from a constant concentration $a$ of actin monomers per unit length, and depolymerization at a rate $k_{\rm d}$. At equilibrium, polymerization and depolymerization balance to give a uniform gel fraction $\phi_0 = a v_{\rm p}/k_{\rm d}$, in terms of which we rewrite the right-hand side in the last equality[2].

#### 2. *Gel force balance*

Force balance in the gel is given by

$$\xi\phi v = \partial_x\sigma. \quad \text{(S2)}$$

The left-hand side corresponds to gel-substrate friction proportional to the coefficient $\xi$ as well as the volume fraction $\phi$ and the velocity $v$ of the gel. Friction arising from cytosol permeation is typically smaller and can be ignored in front of substrate friction[1]. The right-hand side corresponds to internal forces in the gel, which arise from gradients in the gel stress $\sigma$. The stress

$$\sigma = \sigma_{\rm v} + \sigma_{\rm a} - \Pi \quad \text{(S3)}$$

includes contributions from gel viscosity

$$\sigma_{\rm v} = \eta\phi\partial_x v, \quad \text{(S4)}$$

myosin-generated isotropic active stress

$$\sigma_{\rm a} = \zeta\phi, \quad \text{(S5)}$$

with $\zeta > 0$ for contractile stresses, and osmotic pressure $\Pi$. The osmotic pressure can be obtained from the gel's thermodynamics as

$$\Pi = \mu[\phi]\phi - f[\phi] \quad \text{(S6)}$$

based on the chemical potential $\mu$ and free-energy density $f$ given by

$$\mu[\phi] = \frac{\delta F}{\delta\phi}, \qquad F = \int \mathrm{d}^3\boldsymbol{r}\, f[\phi]. \quad \text{(S7)}$$

To obtain these quantities, we take the free energy from the Flory-Huggins theory of polymer solutions[3,4]:

$$F = \int \mathrm{d}^3\boldsymbol{r}\, k_B T\rho \left[\frac{\phi}{N}\ln\phi + (1-\phi)\ln(1-\phi) + \chi\phi(1-\phi) + \frac{\kappa}{2}(\boldsymbol{\nabla}\phi)^2\right]. \quad \text{(S8)}$$

Here, $\rho$ is the three-dimensional monomer concentration, which is directly related to the one-dimensional concentration $a$ introduced in Eq. (S1). The first two terms of Eq. (S8) correspond to the entropy of mixing between the polymer and the solvent, with $N$ the number of monomers. The two last terms account for interactions. The first one corresponds to polymer-solvent repulsion, which can induce passive phase separation[3,4]. As we focus here on phase separation induced by active stresses, we take $\chi = 0$. The last term represents the interfacial energy associated with concentration gradients, with a coefficient $\kappa > 0$. Using Eq. (S6), we obtain the osmotic pressure as

$$\Pi = k_B T\rho\left[\left(\frac{1}{N} - 1\right)\phi - \ln(1-\phi) - \kappa\phi\nabla^2\phi - \frac{\kappa}{2}(\boldsymbol{\nabla}\phi)^2\right]. \quad \text{(S9)}$$

The physical role of the osmotic pressure in this problem is to saturate the activity-driven instability to dictate the amplitude of the final inhomogeneous gel fraction profile $\phi(x-Vt)$. Accordingly, we approximate Eq. (S9) by expanding $\phi = \phi_0 + \delta\phi$ around the equilibrium volume fraction $\phi_0$, and we retain only the third-order term in $\delta\phi$, which saturates the instability[1]. Thus, the osmotic pressure is approximated as

$$\Pi = \alpha(\phi - \phi_0)^3 - \gamma\nabla^2\phi, \quad \text{(S10)}$$

with the coefficients defined as

$$\alpha \equiv \frac{k_B T\rho}{3(1-\phi_0)^3}, \qquad \gamma \equiv k_B T\rho\kappa\phi_0. \quad \text{(S11)}$$

#### 3. Combined equations and boundary conditions

Combining all the contributions, the force balance Eq. (S2) reads

$$\xi v\phi = \partial_x\left[\eta\phi\partial_x v + \zeta\phi - \alpha(\phi-\phi_0)^3 + \gamma\partial_x^2\phi\right]. \quad \text{(S12)}$$

Ignoring the viscous stresses, which are typically smaller than substrate friction[1], we can use this equation to substitute the gel flux $\phi v$ in the mass balance Eq. (S1), which gives

$$\partial_t\phi - V\partial_x\phi + \partial_x\left\{\frac{1}{\xi}\partial_x\left[\zeta\phi - \alpha(\phi-\phi_0)^3 + \gamma\partial_x^2\phi\right]\right\} = -k_{\mathrm{d}}(\phi-\phi_0). \quad \text{(S13)}$$

This equation combines features of the Cahn-Hilliard equation (third term), whereby activity produces phase separation, with a reaction term (right-hand side) that arrests it, and with an advection term (second term) that allows the resulting inhomogeneous steady states to move[1].

To solve Eq. (S13), we specify the following boundary conditions. First, we impose no flux of gel through the cell ends at $x = 0, L$. The gel flux in the cell's frame is $J = \phi(v - V)$, from where

$$v(0) = v(L) = V, \quad \text{(S14)}$$

with $v$ obtained from Eq. (S12). Second, we impose the so-called variational boundary conditions, given by

$$\partial_x\phi|_{x=0} = \partial_x\phi|_{x=L} = 0, \quad \text{(S15)}$$

which assume that the gel has no additional binding energy or dissipation at the cell ends[1].

#### 4. Cell force balance

Equation (S13) contains the cell velocity $V$, which is not a parameter but instead depends on the gel concentration profile, that is, on the solution of Eq. (S13). To determine $V$, we impose a global force balance on the cell:

$$\xi_{\mathrm{f}} V + \int_0^L \xi\phi v\,\mathrm{d}x = 0. \quad \text{(S16)}$$

This equation balances two friction forces: the integrated surface friction discussed in SI Section A 2, which depends on actomyosin flows that provide thrust, and a drag force with coefficient $\xi_{\mathrm{f}}$ that opposes cell motion along the channel. An important contribution to this drag is due to extracellular fluid moving past the cell, either around it in a lubrication layer or through it via macropinocytosis[5]. Therefore, this fluid drag exists even without substrate friction, for example if the cell is externally pulled along a channel with frictionless walls.

Together, Eqs. (S13) and (S16) with the boundary conditions Eqs. (S14) and (S15) provide a minimal self-consistent set of equations to describe amoeboid migration.

### B. Frictiotaxis

To study the mechanism of adhesion-independent durotaxis, we consider the model introduced in SI Section A in the presence of inhomogeneous friction $\xi(x)$. As explained in the Main Text, we suggest a mechanism whereby a friction gradient breaks the symmetry and polarizes cells up the gradient.

As the cell poles are not subject to substrate friction (Fig. 5b), they undergo the contractile instability of Ref.[1] at a faster rate than the rest of the cortical gel. As a result, contractile actomyosin foci appear at or near the cell poles. The flows generated by pole contraction produce a stress $\sigma_{\mathrm{pole}}$ that pulls on the neighboring regions of the cortex. In particular, the pole stress pulls on the central region of the cell, in which friction with the substrate makes the contractile instability slower. Hence, we consider the central region to start in the unpolarized state with uniform gel concentration $\phi_0$, for which the force balance Eq. (S12) reduces to

$$\xi v = \eta\frac{\mathrm{d}^2 v}{\mathrm{d}x^2}, \quad \text{(S17)}$$

as given in Eq. 2 in the Main Text.

To illustrate the proposed mechanism of symmetry breaking, we obtain the cortical flow induced by pole contraction at the ends of the central cell region, located at $x = \pm L/2$. For simplicity, we solve Eq. (S17) by taking a different constant friction coefficient for each end, $\xi_-$ and $\xi_+$, with $\xi_- < \xi_+$. Imposing $\sigma(\pm L/2) = \sigma_{\mathrm{pole}}$, the solution of Eq. (S17) around each end is

$$v_+(x) = \frac{\sigma_{\mathrm{pole}}\lambda_+}{\eta}e^{(x-L/2)/\lambda_+}, \quad \text{(S18a)}$$

$$v_-(x) = -\frac{\sigma_{\mathrm{pole}}\lambda_-}{\eta}e^{-(x+L/2)/\lambda_-}, \quad \text{(S18b)}$$

where $\lambda_\pm \equiv \sqrt{\eta/\xi_\pm}$ is the hydrodynamic screening length that determines the decay length of the flow field. Because $\xi_+ > \xi_-$, we have $\lambda_+ < \lambda_-$, and hence the flow field decays over a longer distance on the low-friction side. The speed is also higher on the low-friction side, with

$$v_\pm \equiv v(\pm L/2) = \pm\frac{\sigma_{\mathrm{pole}}\lambda_\pm}{\eta} = \pm\frac{\sigma_{\mathrm{pole}}}{\sqrt{\eta\xi_\pm}} \quad \text{(S19)}$$

as quoted in Eq. 3 in the Main Text. By virtue of the advective term in Eq. (S1), the higher speed at the low-friction side will produce faster increase in gel concentration on that side, thus breaking the symmetry of the cortex as illustrated in Fig. 5b.

1. Callan-Jones, A. C. & Voituriez, R. Active gel model of amoeboid cell motility. *New J. Phys.* **15**, 025022 (2013).
2. Hawkins, R. J. *et al.* Spontaneous Contractility-Mediated Cortical Flow Generates Cell Migration in Three-Dimensional Environments. *Biophys. J.* **101**, 1041–1045 (2011).
3. Rubinstein, M. & Colby, R. H. *Polymer Physics* (Oxford University Press, Oxford, 2003).
4. Doi, M. *Introduction to Polymer Physics* (Oxford University Press, Oxford, 1995).
5. Bergert, M. *et al.* Force transmission during adhesion-independent migration. *Nat. Cell Biol.* **17**, 524–529 (2015).
